# Enhancement of Protein Stability by Quenching Millisecond Conformational Dynamics

**DOI:** 10.1101/2023.08.30.555640

**Authors:** Xue-Ni Hou, Chang Zhao, Bin Song, Mei-Xia Ruan, Xu Dong, Zhou Gong, Yu-Xiang Weng, Jie Zheng, Chun Tang

## Abstract

Protein folding may involve folding intermediates. Ubiquitin (Ub) is a 76-residue small protein essential in post-translational modification and cell signaling. Ub is also a model system of protein folding. Previous studies have indicated the involvement of a folding intermediate, as Ub C-terminal residues, including strand β5, only dock correctly at a later stage. The natively folded Ub undergoes conformational dynamics over a vast range of timescales. At the millisecond timescale, Ub transiently digresses to a C-terminally retracted state, which is extremely rare and has only been recently identified at an elevated temperature. Herein through a conjoint use of NMR, MS, and MD simulations, we have established a link between Ub millisecond dynamics and protein stability. Among the alanine mutations that have been systematically introduced to the hydrophobic residues in β5, L67A and L69A elevate the population of the retracted state and enhance conformational interconversion, which facilitates the undocking of β5 and the exposure of protein hydrophobic core. Conversely, L71A and L73A mutations decrease the population of the retracted state and quench millisecond dynamics, which causes a significant enhancement of protein stability. As such, the transition state of Ub millisecond dynamics is the much sought-after folding intermediate, whereas C-terminal mutations alleviate the dependence on this intermediate and reduce the unfolding to an all-or-none process. Though having a negative impact on protein stability, Ub millisecond dynamics likely facilitate proper protein turnover and allow the fulfillment of its biological function.

## Introduction

Most proteins fold into their native structure at microsecond to millisecond timescale (1–4). Many single-domain proteins fold rapidly upon the collapse of the hydrophobic core, involving only two states—an unfolded state and a natively folded state (5–7). Alternatively, one or more folding intermediates can be involved, either on-pathway or off-pathway (4, 8–20). When folded into the native conformation, a protein undergoes hierarchical dynamic fluctuations over a vast range of timescales, thus to fulfill specific biological functions (21). Studies have elucidated the structures of lowly-populated on-pathway folding intermediates that rapidly exchange with the native protein conformations (19, 20, 22). However, it is unknown whether folding intermediates exist in the two-state all-or-none folding processes, albeit they have eluded direct characterization due to extremely low populations. It is also unclear how protein dynamics—the transient digression to an alternative conformation—is related to protein folding, which may represent an off-pathway folding intermediate and serve as a kinetic bump to the folding/unfolding process. A more pertinent question is how protein dynamics involving the folding intermediates impact protein stability.

Ubiquitin (Ub), a 76-residue single-domain protein, is involved in myriad aspects of cell signaling. The primary function of Ub is to covalently modify a cellular protein via an isopeptide bond, and to target the modified protein for proteasomal degradation (23–26). Ub has five β-strands––β1 through β5––all located on the same side of the protein. In addition, two α-helices, α1 and α2, are located after β2 and before β5, respectively. The folding mechanism of Ub has been extensively investigated (4). It has been shown with denaturant-jump experiments that Ub folds into its native conformation at a low-millisecond timescale (6). However, hydrogen-deuterium exchange and protection analysis have shown that the C-terminal residues involving β5 and the preceding α2 likely fold at a late stage and take much longer than the rest of the protein to assume the correct conformation (27, 28). Mutational scanning and Φ-value analysis have further indicated that an intermediate state is likely involved, and the Ub folding process at an ambient temperature can be characterized by a three-state process (9–11, 29). Nevertheless, experimental investigations of Ub folding/unfolding fell short in providing structural details for the intermediate state. Moreover, a two-state folding model seems to suffice to describe the Ub folding process (15, 29, 30).

Ub is highly dynamic. NMR studies have shown that Ub fluctuates at the μs-ms timescale, with the β-strands dynamically curling up (31–33). The β-strands form the primary binding surface for Ub partner proteins, as the conformational dynamics at low-μs timescale are responsible for recognition specificity. In 2015, Komander and his coworkers showed that phosphorylation by the kinase PINK1 at Ub residue S65 elicits an alternative conformation, namely the C-terminally retracted state, or simply the retracted state (34). The retracted state of the phosphorylated Ub (pUb) mainly differs from the C-terminally relaxed state (i.e., the native Ub fold) in the hydrogen bond pattern of β5, which moves up by two residues (35). Under the physiological conditions, the pUb retracted state is about equally populated to the relaxed state, but increases with pH (35).

It turns out that the unmodified Ub already undergoes an interconversion between relaxed and retracted states, though the latter has long eluded detection. The retracted state of the wildtype Ub is extremely lowly populated, and could only be detected at an elevated temperature (45 ºC, 318 K) using the sensitive NMR chemical exchange saturation transfer (CEST) experiment (36). Of note, CEST NMR analysis shows that the interconversion between the two Ub conformational states occurs at a millisecond timescale (37). Phosphorylation at Ub residue S65 thus can be considered a special mutation that destabilizes the native state and enhances the population of the retracted state. Indeed, the S65 phosphoryl group is partially and unfavorably buried in a hydrophobic patch, as manifested by its unusually large pKa value (35).

These previous studies prompted us to evaluate the relationship between Ub millisecond dynamics and its folding/unfolding process, with special attention to the correct positioning of β5. We systematically introduced alanine mutations to the buried hydrophobic residues in Ub β5 (Figure 1), and showed that the mutations either promote the conversion to the retracted state (L67A), enhance the interconversion rate to the retracted state (L69A), or quench the interconversion (L71A and L73A). Significantly, L71A and L73A mutations enhance Ub thermal stability so much that the L73A mutant has a melting temperature (*T*_m_) exceeding 100 ºC.

**Figure 1.**
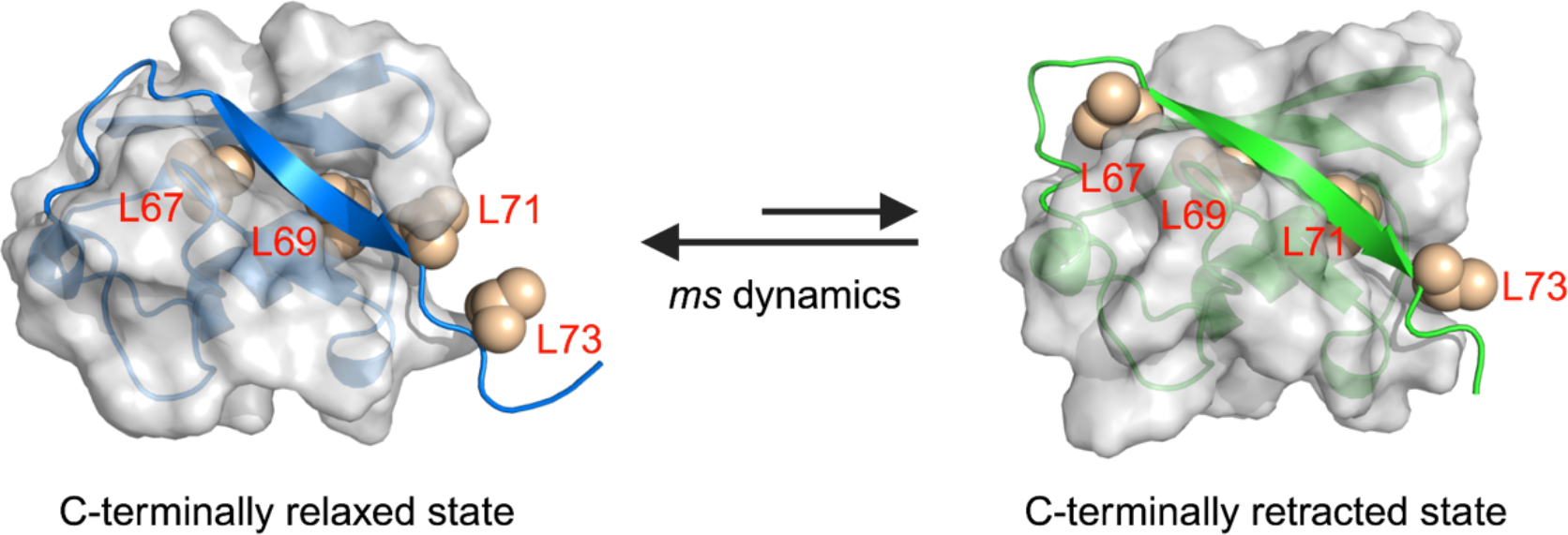
Ub adopts two alternative conformations. A schematic showing the structural difference between the C-terminally relaxed and retracted states. In the latter structure, β5 moves up by two residues, thus burying a different set of hydrophobic residues, as indicated with spheres. The retracted state structure is generated by removing the phosphoryl group from the known pUb structure, which is then equilibrated with MD simulation.

## Results

### Point mutations in β5 alter Ub thermal stability

We introduced alanine mutations, one at a time, to Ub residues L67, L69, L71, or L73. These residues are located in β5, pointing inward. In the native state, the side chains of residues L67 and L69 are buried in the protein hydrophobic core, residue L71 is partially buried, and residue L73 is located in the C-terminal flexible tail and fully exposed. Previous structural characterization of the pUb has shown that, in the retracted state, residues L69 and L71 are fully buried, while residues L67 and L73 are partially buried (Figure 1B).

We used differential scanning fluorometry (DSF) of the intrinsic tyrosine fluorescence to assess the thermal stability of the wildtype, mutant, as well as phosphorylated Ub proteins (38). We evaluated the temperature-dependent change of the relative fluorescence at 330 nm and 350 nm — the *T*_m_ value can be determined by taking the first derivative of the fluorescence ratio. Consistent with the previous report (37), phosphorylation at residue S65 decreases the *T*_m_ value of Ub by more than 10 ºC. The L67A and L69A mutants also have lower *T*_m_ values than the wildtype, but higher than the pUb. In contrast, the L71A and L73A mutations elevate the *T*_m_ value. Significantly, the *T*_m_ value of the L73A mutant is about 100 ºC (Figure 2A-D). Note that the DSF derivative curve of the L73A mutant shows a broad peak, likely due to the unusually high melting temperature.

**Figure 2.**
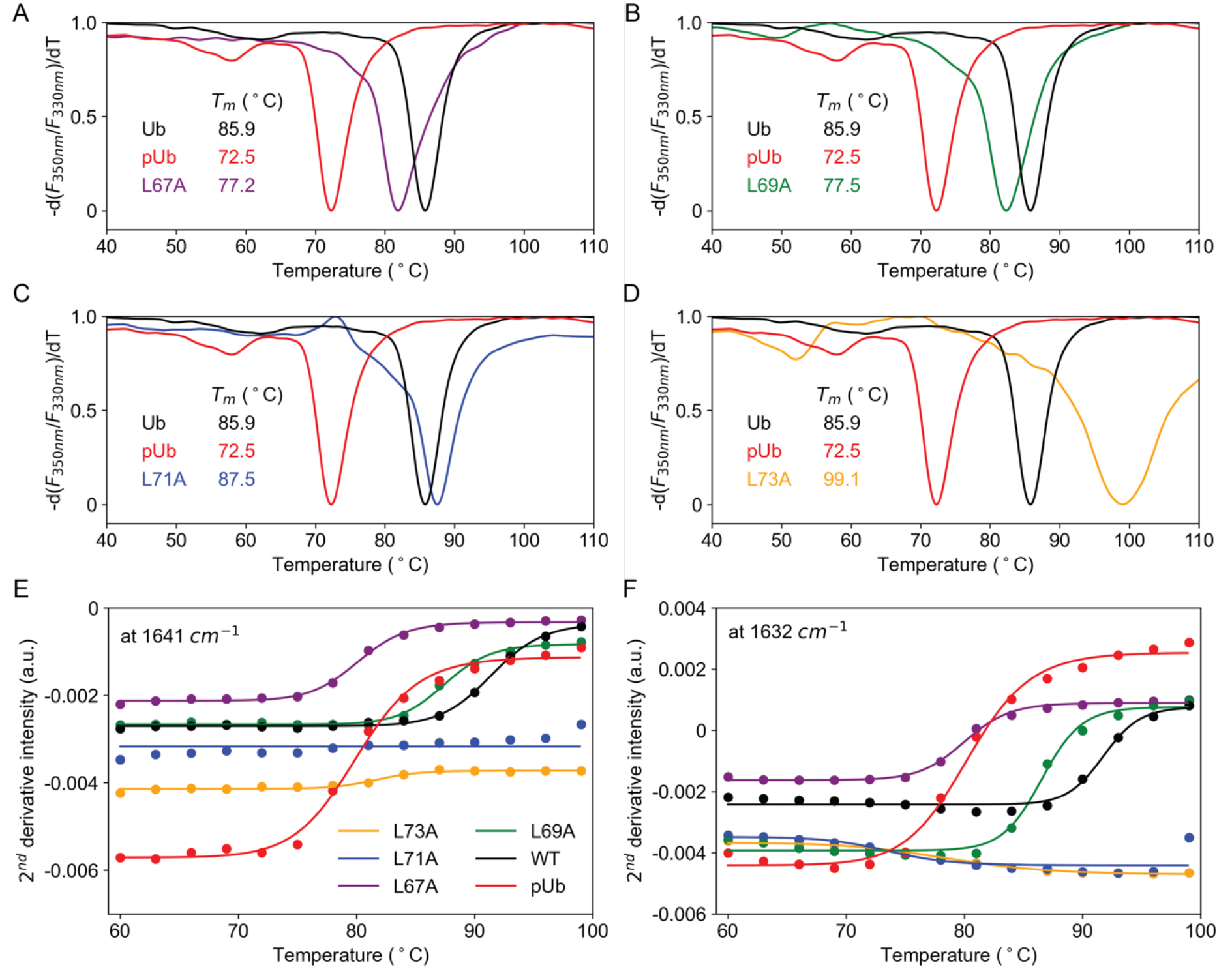
Thermal stability of Ub proteins assessed with DSF and FTIR. (A-D) First derivatives of the ratios of fluorescence intensities at 350 nm and 330 nm for L67A (A), L69A (B), L71A (C), and L73A (D), respectively. The curves for Ub wildtype (black line) and phosphorated Ub (red line) are plotted for comparison. (E, F) Second derivatives of FTIR thermal titration curves in D_2_O of various Ub constructs at 1641 cm ^-1^ (E) and 1632 cm ^-1^ (F). For clarity, the data were tentatively fitted to solid sigmoid curves.

We also assessed the thermal stability of the Ub mutants using Fourier-transformed infrared (FTIR) spectroscopy. We collected a series of FTIR spectra for the various versions of Ub proteins at increasing temperatures. For the wildtype Ub, the changes in the vibrational spectra are visible at the temperature above 87 ºC, as characterized by a decrease in intensity of the 1641 cm^-1^ band and a concomitant increase of intensity of the 1620 cm^-1^ band (Figure S1). The amide-I band at 1641 cm^-1^ results from the collective vibration of the Ub anti-parallel β-sheet, while the 1620 cm^-1^ band results from the vibrational mode for the unstructured protein (12, 39). Thus, the changes in the FTIR absorbance and its second derivative (Figure 2E) are correlated with the loss of secondary structure and overall protein heat denaturation. The changes in the FTIR spectra occur at much lower temperatures for the L67A and L69A mutants and pUb than Ub wildtype (Figure 2E). In contrast, the FTIR spectra of the L71A mutant only start to change at the highest temperatures assessed, whereas the FTIR spectra of the L73A mutant display little changes across the entire temperature range (Figure 2F). Taken together, alanine mutations to the C-terminal leucine residues result in an enhancement of Ub thermal stability.

### L67A mutation promotes the conversion to the retracted state

To understand the underlying cause for the altered thermal stability of the mutants, we resorted to NMR spectroscopy and characterized how the point mutations affect Ub structure and dynamics. The ^1^H-^15^N heteronuclear single-quantum coherence (HSQC) spectra of the L73A mutant is similar to that of the wildtype Ub, except for the residues immediately next to the mutated residue (Figure S2). The L69A and L71A mutations caused small perturbations to the HSQC spectrum, which also include nearby residues in β1 and β3 (Figures S3 and S4). Interestingly, the HSQC spectrum of the Ub L67A mutant displays more peaks than the number of non-proline residues, indicating the presence of multiple conformational states. Indeed, two sets of peaks with distinct intensities can be identified and assigned (Figure S5A).

The peaks of the L67A mutant can be assigned to two different species. Based on the relative peak intensities in the NMR spectrum, the major and minor species are populated at about 82% and 18%, respectively, at 310 K. Extensive chemical shift differences are observed between the major species and the wildtype Ub, involving almost all β-sheet residues, whereas much smaller chemical shift differences are observed for the minor species of L67A mutant (Figure 3A-B). It has been shown that mutation of Ub residue L67 to a polar residue, i.e., serine or asparagine, causes a full conversion to the retracted state (37). Here L67A mutation also promotes the conversion to the retracted state, albeit incompletely, which constitutes the major species. Indeed, the Cα chemical shift values of residues R72 and L73 of the major species of L67A mutant are characteristic of the secondary shift of β-sheet structure (Table S1).

**Figure 3.**
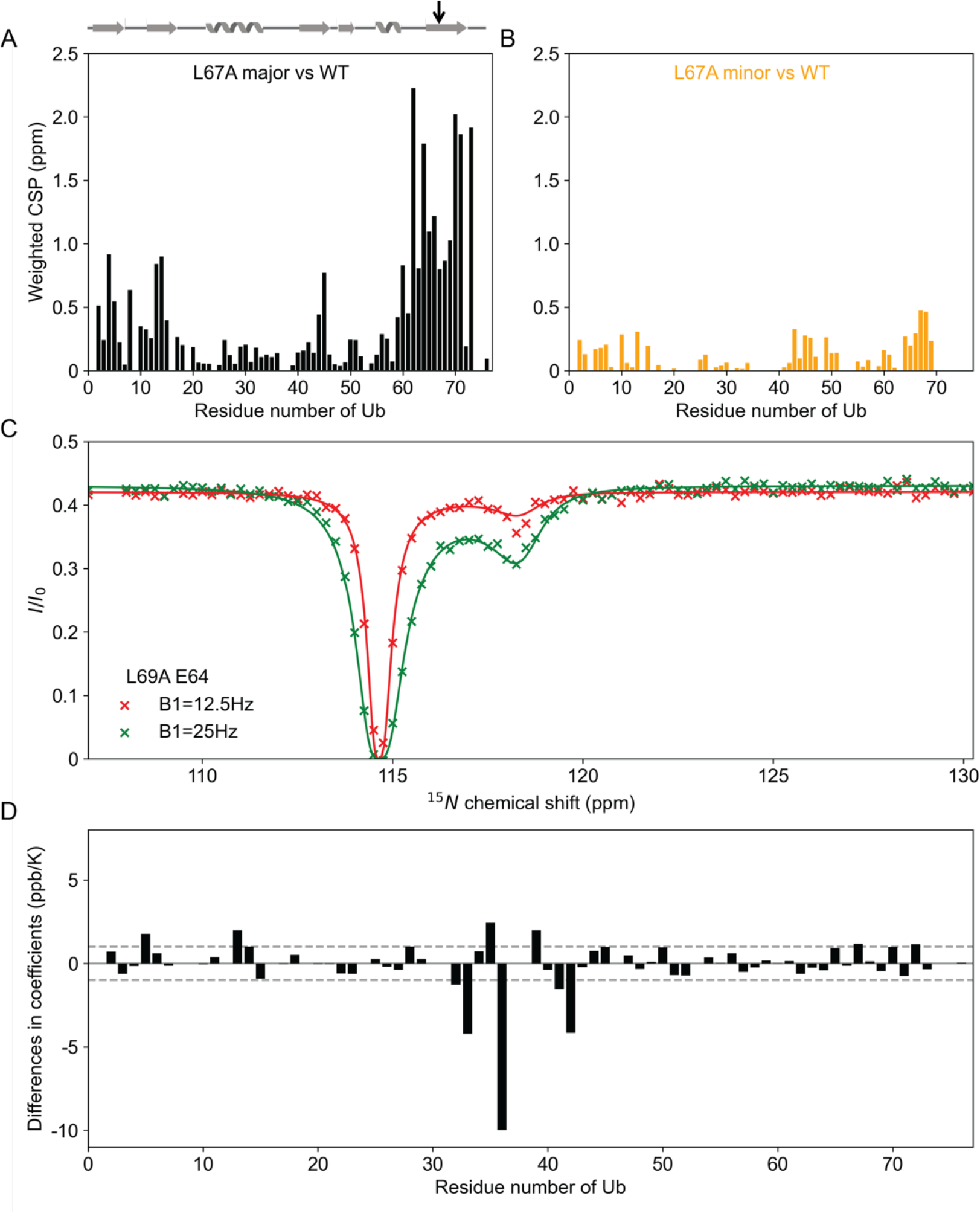
L67A and L69A mutations cause perturbations to the structure and dynamics of Ub as monitored by NMR. (A, B) Chemical shift perturbations (CSPs) between major state (A) and minor state (B) of Ub L67A mutant from the Ub wildtype. Ub secondary structure elements and the mutated residue is denoted at the top of the panel (A). The point mutation introduced is indicated by an arrow. (C) L69A mutant undergoes pronounced millisecond dynamics. Representative peak fits for CEST experiments on Ub L69A. The exchange between major and minor species is determined at ∼360 s^-1^, and the minor species (retracted state) is determined at a population of 1.3 %. (D) Differences in amide proton temperature coefficients between Ub L69A mutant and wildtype.

The coexistence of the two conformational states of the Ub L67A mutant allowed us to assess the thermodynamics of their conversion based on the NMR peak intensities. The retracted state amounts to ∼83% at 318 K, but decreases to ∼72% at 286 K (Figure S5B). Thus, the unimolecular conversion from Ub relaxed state to the retracted state is endothermic, which has also been shown to be the case for the pUb (40).

### L69A mutation accelerates Ub millisecond dynamics

A point mutation in β5 not only changes the relative population of the two conformational states of Ub, and also changes the interconversion timescale. Using CEST NMR, we assessed the overall exchange rate *k*_ex_, i.e., the sum of forward and reverse exchange rates, between the relaxed state and retracted state of Ub L67A mutant at two *B*1 fields at 318 K, affording 29.6 ± 7.0 s^-1^ (Figure S6). The exchange rate is consistent with the fact that two sets of peaks are resolved in the HSQC spectrum for the L67A mutant, and is close to the value for the wildtype Ub at 55.1±14.1 s^-1^ (Figure S7).

Using CEST NMR, we assessed the millisecond dynamics of the Ub L69A mutant. We determined that the retracted state is populated 1.30 ± 0.03% at 318K, higher than that of the wildtype Ub. Significantly, the determined interconversion rate *k*_ex_ between relaxed and retracted states is 360.2±20.0 s^-1^, almost an order of magnitude faster than that of the wildtype (Figure 3C). Thus, the energy barrier between the two Ub conformational states is lowered when introducing the L69A mutation.

We also assessed the ^1^H-^15^N chemical shift values over a range of temperatures (Figure S8). A temperature coefficient more negative than –4.5 ppb/Kelvin indicates the amide proton is unprotected by hydrogen bonding. The temperature coefficients for the wildtype Ub are consistent with the previous reports (41, 42). The L69A mutant has more labile residues, with residues K33, I36, and R42 displaying more negative temperature coefficient of their amide protons (Figure S9 and Figure 3D). The results are consistent with the CEST NMR analysis and further indicate the L69A mutant undergoes more pronounced millisecond dynamics.

More evidence of the structural destabilization upon the point mutations in relationship to protein dynamics can be obtained from hydrogen-deuterium exchange analysis by mass spectrometry (HDX-MS) (43, 44). We assessed the time-dependent deuterium exchange rates for the peptides derived from the mutant and wildtype Ub proteins at 323 K. Solvent exchange rates for the C-terminal peptide are faster for the L69A mutant than for the L67A mutant, confirming that the transition state, presumably characterized with a solvent-exposed fully open β5, can be more readily accessible by the L69A mutant (Figure S10 and Figure 4). Note that in an EX2 scheme of HDX, a structural element has to first open up, with hydrogen bonds of the native conformational state broken, in order to allow the solvent exchange to take place (43). Thus, the open state for deuterium exchange likely represents the transitional state of the interconversion between the relaxed and retracted states.

**Figure 4.**
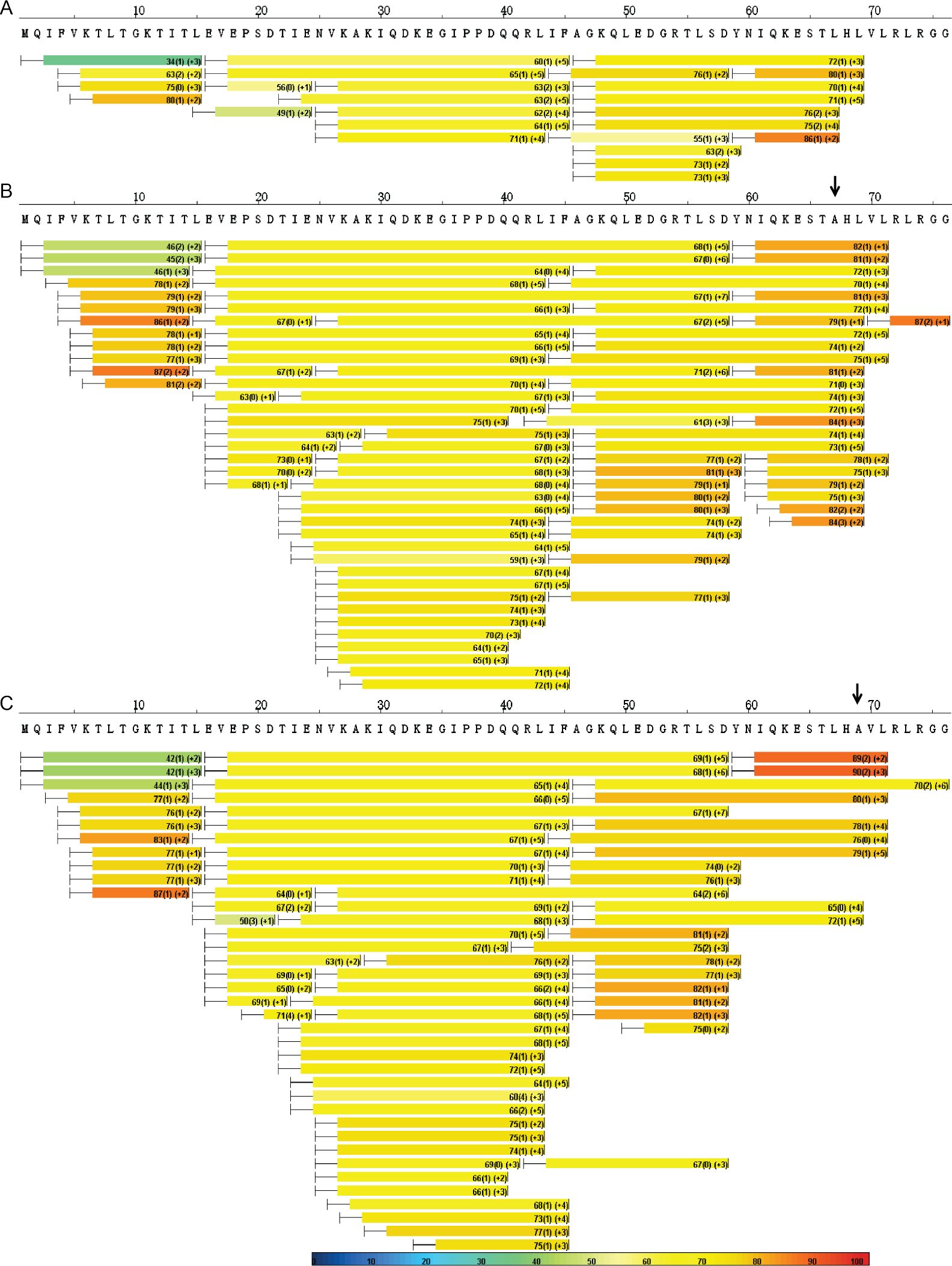
L67A and L69A mutations promote the structural opening and solvent exchange of Ub. Heat maps of HDX-MS analyses for the wildtype (A), L67A (B), and L69A (C) Ub proteins at 323 K upon D_2_O exchange for 45 s. The level of deuterium substitution ratio of the labile protons is color-coded, as indicated by the legend at the bottom. The point mutation introduced is indicated by an arrow.

On the other hand, the exchanging rates for the N-terminal peptides derived from L67A and L69A Ub mutants carrying various charges are faster than the equivalent peptides derived from the wildtype protein (Figure S11). This indicates that the point mutations not only facilitate the exposure and undocking of the β5 but also loosen up the protein hydrophobic core.

We obtained few exchange data for the C-terminal residues of the wildtype Ub mutant, which is likely resistant to proteolysis at the experimental temperature (Figure S10). However, increasing the temperature from 323 K to 343 K could result in more exchange (Figure S12), confirming a temperature-accelerated exchange process by overcoming the activation energy of opening up the β5.

### L71A and L73A mutations quench Ub millisecond dynamics

We also performed CEST measurements for Ub L71A and L73A mutants, but observed no minor dip in the CEST profiles. This means that the two mutations negatively impact the millisecond dynamics of Ub. Further evidence for the quenched millisecond dynamics comes from the phosphorylation assay. It has been shown that the PINK1 kinase catalyzes the phosphorylation of Ub only in its retracted state, in which residue S65 becomes exposed and fits into the active site of the kinase (45). The phosphorylation rates are drastically slowed for both L71A and L73A mutants, in comparison to the wildtype Ub and L69A mutant. No phosphorylated species could be detected for the L71A or L73A mutant after more than an hour, and could only be observed after more than three hours of phosphorylation reaction (Figure S13).

Despite being a poorer substrate for PINK1 kinase, the L73A mutant still binds to its partner proteins efficiently. Titrating the UBA domain (residues 578-621 from ubiquilin-2, ref. (46)), we determined the binding affinities toward the wildtype Ub and L73A mutant are close (Figure S14). Thus, the ns-μs dynamics of Ub, which involve the curling movement of the β-sheet and are responsible for protein association (47, 48), seem unaffected.

### Energetic perturbation of Ub point mutations

To gain additional mechanistic insight, we resorted to molecular dynamics (MD) simulations and assessed the change of Ub dynamics upon the mutations. From the unrestrained MD simulation trajectories of the L71A and L73A mutants, we observed the fluctuations of the distance between β5 and the rest of the Ub protein are no different from that of the wildtype Ub (Figure S15). In comparison, the relative position of β5 undergoes larger fluctuations for the L69A mutant than for the wildtype, averaged at 7.41±0.64 Å and 7.15±0.37 Å, respectively (Figure 5A).

**Figure 5.**
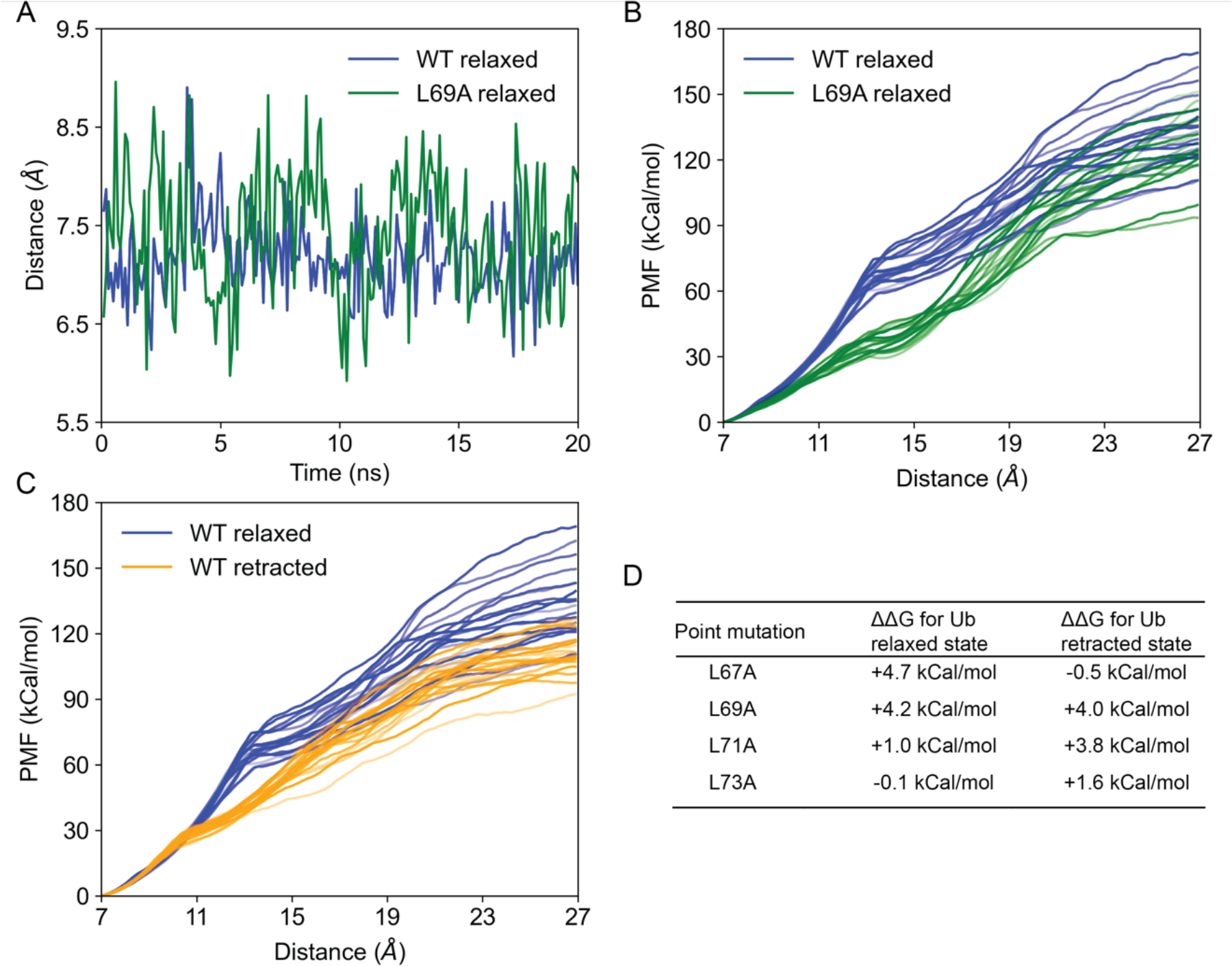
Computational analysis of wildtype and mutant Ub proteins at 318K. (A) Unrestrained MD simulations for Ub wildtype and L69A mutant, both in relax-state conformation. The mutant exhibits a larger fluctuation in the relative position of β5, as indicated by the Cβ distance between residues L30 and I30, 7.41±0.64 Å vs. 7.15±0.37 Å. (B, C) Steered MD simulations for Ub wildtype in the relaxed state, L69A mutant in the relaxed state, and Ub wildtype in the retracted state. A constant force is applied at the L69 Cβ atom of the relaxed state or the L71 Cβ atom of the retracted state. 20 independent trajectories are plotted. (D) Destabilization energy DDG calculated for each mutant as referenced to either wildtype Ub relaxed state or retracted state.

We then performed steered MD (SMD) simulation to recapitulate the β5-detached transition state by pulling the β5 away. Less work is required to pull the β5 out for the L69A mutant than for the wildtype. Moreover, a sharp transition at a distance of ∼13 Å is observed in the energy profile of the wildtype, whereas the SMD energy increases rather gradually for the L69A mutant (Figure 5B). On the other hand, the SMD trajectories of the wildtype Ub in the retracted state also lack a sharp transition and require less work to open up the β5 (Figure 5C). Together, the relaxed state of Ub wildtype has the most cooperative hydrogen bonds for unfolding, which is consistent with both NMR and MS results.

For the L69A mutant, it appears that the increased dynamics at a slower timescale, evidenced by the lower work required to pull open the β5, is hierarchically coupled to the increased dynamics at a fast timescale, evidenced by the unrestrained MD simulations (Figure 5A). The unrestrained MD simulations of Ub wildtype show a smaller fluctuation of the β5 position at 288 K than at 318 K (Figure S16), which partly explains that more work is required to pull open the β5 at the low temperature (Figure S16C). Thus, the β5 is less likely detached, and the transition state is less accessible at a low temperature.

Evaluation of the change in free energy upon the point mutation further establishes a link between protein dynamics and stability (Figure 5D). L67A mutation destabilizes the relaxed state by about 4.7 kCal/mol, but has almost no effect for the retracted state. L69A mutation, on the other hand, destabilizes the relaxed and retracted states by a similar amount, thus narrowing the energy gap between the relaxed/retracted states and the transition state. Thus, for the L69A mutant, lower activation energy is required for the conformational interconversion, and the transition state is more accessible at the same temperature. On the other hand, the L71A and L73A mutations destabilize the retracted state, but have little to no effect on the energetics of the relaxed state.

## Discussion

The folding of Ub occurs at the millisecond timescale. Previous studies have indicated that Ub folding proceeds via a three-state pathway at ambient temperature, as the correct positioning of the C-terminal residues takes much longer than the rest of the protein (27, 28). It has also been noted that a two-state folding mechanism at a relatively low temperature seems to suffice (15, 29, 30). On the other hand, the natively folded Ub dynamically interconverts between two alternative conformational states, also at the millisecond timescale. In the present study, we systematically characterized Ub alanine mutants with an integrative use of NMR, HDX-MS, and MD simulations, and showed that the transition state between Ub relaxed and retracted state is likely the long-sought-after Ub folding intermediate. We characterized Ub dynamics as high as 45 ºC, 318 K, a temperature very taxing for the NMR instrument. Thanks to the conjoint use of HDX-MS, we could characterize the protein dynamics at higher temperatures. Moreover, MD simulations provided additional mechanistic details.

The effect of substituting a nonpolar residue with a less hydrophobic or polar residue on protein stability has been well-characterized (49). In a previous study, introducing an L67A mutation has also been shown to destabilize Ub by a similar amount (50). But surprisingly, both L71A and L73A mutants are more thermally stable than the wildtype, as shown with the FTIR and DSF measurements. The L73A mutant becomes so stable that it remains folded at a temperature >100 ºC. Accordingly, we found that the millisecond dynamics between Ub relaxed and retracted states are quenched upon the mutation, as shown with NMR CEST analysis and phosphorylation assay. The increased stability of these two mutants results from the destabilization of the retracted state. Consequently, Ub unfolding involving the β5-open transition state is less opted (Figure 6).

**Figure 6.**
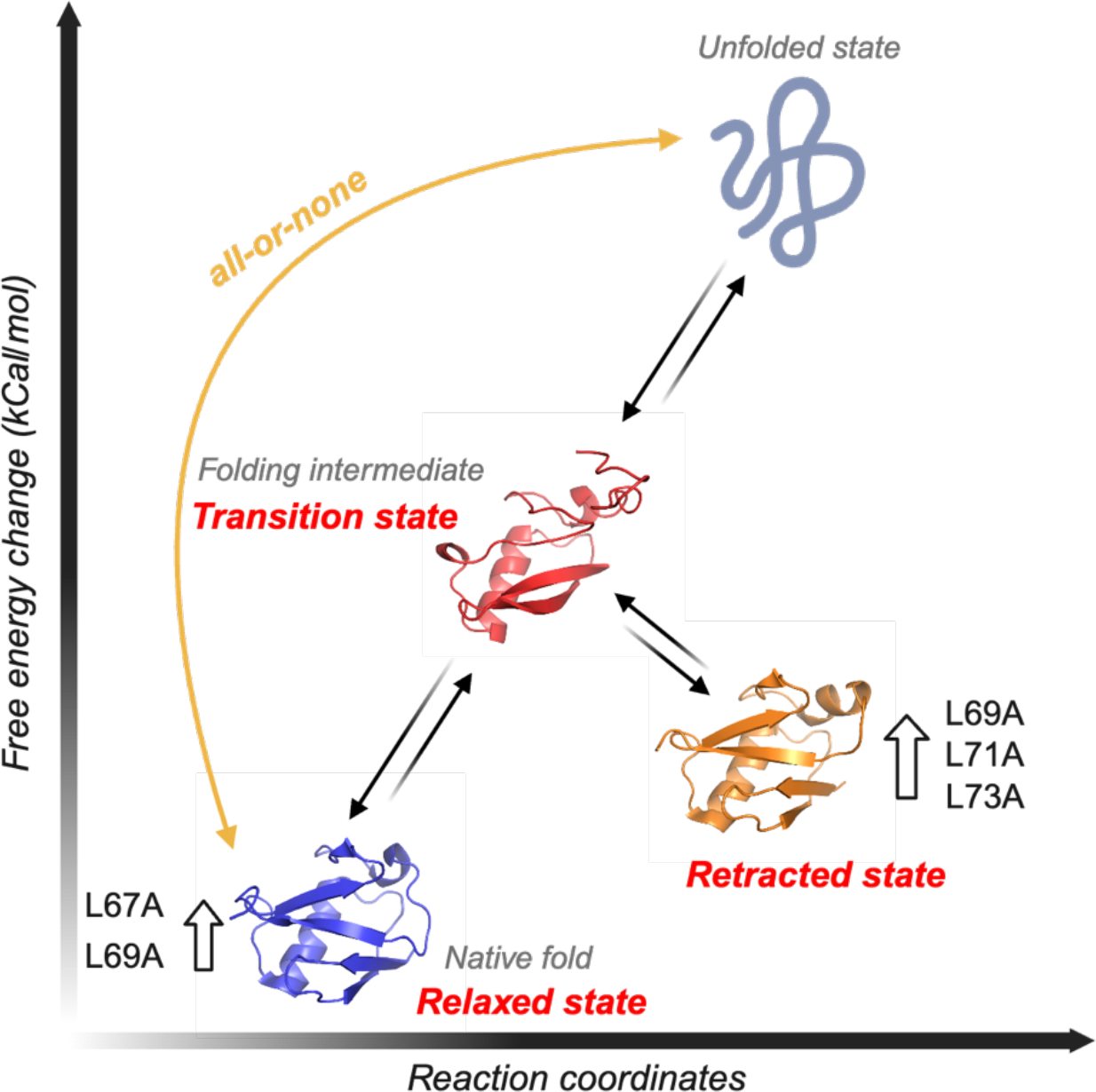
A schematic showing how the folding intermediate destabilizes Ub. As presented, L67A mutation destabilized Ub relaxed state, i.e., the native fold; L69A mutation destabilizes both relaxed and retracted states, which makes the transition state more accessible and accelerates millisecond dynamics; L71A and L73A mutations destabilize the retracted state and makes the interconversion unfavorable. As such, decreasing the involvement of the intermediate pathway can reduce the unfolding to an all-or-none process and thus enhance protein stability.

Per microscopic reversibility, the transition state between Ub relaxed and retracted states can be equated to Ub folding intermediate. However, the β5-undocked intermediate occupies a very high energy level and extremely low population, preventing direct characterization (Figure 6). Such energy landscape has to do with the high stability of the wildtype Ub, which has a melting temperature of >80 ºC (Figure 2). Therefore, the folding towards Ub native conformation is a highly energetic favorable process. Notwithstanding, this folding intermediate is likely a flaw designed to make Ub more dynamic and less stable.

The C-terminally retracted state represents an off-pathway intermediate of Ub folding. Here the C-terminal β5 is positioned differently, resulting in different hydrogen bond network, and consequently, weaker cooperativity. Owing to the dynamic interconversion between the two conformational states at the millisecond timescale, Ub can unfold more readily through this transition state, an alternative and likely more efficient pathway towards the unfolded state than the all-or-none pathway (Figure 6). Introducing L71A or L73A mutation decreases the population of the retracted state and quenches the interconversion dynamics, thus reducing the flux through this alternative unfolding pathway. On the other hand, mutations to β5 residues should have little effect on the energetics of the transition state ensemble, as shown previously with the Φ-value analysis (4), and therefore, the folding rate is likely unaffected. Moreover, since the conversion from Ub relaxed state to the retracted state is endothermic, the occupancy of the retracted state would decrease at low temperatures, and folding/unfolding would reduce to a two-state process (15, 29, 30).

Other than being the preferred substrate of PINK1 (45, 51), the biological function of the Ub retracted state remains to be fully established. Yet, Nature may have intentionally incorporated this late-folding intermediate towards Ub native structure. When targeting ubiquitinated proteins for proteasomal degradation, some Ub moieties in the Ub chain, particularly the one immediately adjacent to the modified proteins, are also hydrolyzed (52–54). Herein, Ub C-terminus is covalently attached to the protein that is about to be degraded—a relatively attainable undocking of the β5 will allow Ub to be threaded through the narrow passage in the proteasome for complete unfolding and degradation (55). Thus, having a super-stable Ub is harmful to proteasomal activity and cell survival. Indeed, mutagenesis screenings have shown that replacing L73 with alanine or other polar residue reduces yeast fitness (56, 57).

In summary, our current study has demonstrated that protein dynamics, in particular those occurring at the millisecond timescale, are coupled with protein folding and have a negative impact on protein stability. Thus, eliminating the folding intermediates and reducing unfolding to a two-state process can result in an enhancement of protein stability.

## Materials and Methods

### Sample Preparation

Wildtype human ubiquitin (Ub) in the pET11a expression vector was modified using the KOD-Plus-Mutagenesis Kit (TOYOBO, Tokyo, Japan) to introduce L67A, L69A, L71A or L73A mutation. Ub, Ub mutants, *pediculus humanus corporis Ph*PINK1 (residues 115 to 575) and human ubiquilin2 UBA domain (residues 578 to 621) were expressed and purified as previously described (34, 46, 58). In this study, the *Ph*PINk1 was further purified by Superdex™ 75pg Increase 10/300GL column (Cytiva) preequilibrated with NMR buffer consisting of 20 mM HEPES at pH 7.4 and 100 mM NaCl following the cleavage of GST-tag by using TEV protease.

### Determination of the Thermal Stability of Ub mutants

The assessment was performed using two different techniques. (1) *Intrinsic Differential Scanning Fluorometry (DSF)*. A Prometheus NT.48 (NanoTemper Technologies GmbH, Munich, Germany) was used to assess Ub melting temperature. The experiment measures the intrinsic fluorescence intensity at 330 and 350 nm with the excitation at 280 nm of the intrinsic fluorescence. Three consecutive temperature scans of the same sample (3 × 10 μL) from 40 to 110 ºC in a linear ramp of 1 ºC/min were performed, with the sample equilibrated for 1 min at each temperature point prior to the measurement. The Ub samples were filled into the standard iDSF (DSF with intrinsic fluorescence) grade capillaries, and all samples were formulated in the same buffer as used for NMR. A negative control with only the NMR buffer was included in each run. The apparent Ub melting temperature (T_m_) was determined as the lowest point in the temperature-dependent variation (first derivative) of the 350nm/330nm fluorescence ratio, as evaluated using the ThermControl software V2.1 (NanoTemper Technologies).

(2) *Fourier-transformed infrared spectroscopy (FTIR)*. FTIR absorption spectra were collected using a Vertex 70V (Bruker Optics, Germany). Ub (wildtype or mutant) solution (3 μL), at a concentration of 10 mg/mL, was placed in a two-compartment CaF_2_ sample cell with a 50 μm thick Teflon spacer; the D_2_O reference was measured simultaneously. The measurements were performed in a home-built vacuum chamber with a temperature controlled at an accuracy of ±0.1 ºC by water circulation (59, 60). The samples were thermally equilibrated for 4 min prior to each measurement, from room temperature to over 100 ºC. Note that the chamber holding the L73A mutant remains clear after reaching the highest temperature. A second derivative analysis was also performed for the FTIR spectra.

### NMR spectroscopy

Nuclear magnetic resonance (NMR) acquisition was performed at 298 K on Bruker Avance III HD 800 MHz spectrometers equipped with cryogenic triple-resonance TCI probes unless otherwise stated. Topspin (Bruker) and NMRPipe were used for data processing, and Poky was used for data analysis (61, 62).

Backbone chemical shift assignments were obtained using standard triple-resonance pulse sequences. HNCACB and HN(CA)CO spectra were collected with 2048 × 40 × 100 complex points in the ^1^H, ^15^N, and ^13^C dimensions, respectively. HN(CA)CO spectra were collected using a non-uniform sampling scheme at a rate of 30% complex points in the ^1^H, ^15^N, and ^13^C dimensions, respectively, and were reconstructed using Iterative Shrinkage Thresholding (IST) in the NMRPipe package (63). Due to the small chemical shift perturbations, cross-peak assignment of L71A and L73A mutants were obtained by a comparison to ^1^H, ^15^N-HSQC spectrum of the wildtype. Upon backbone assignment, weighted chemical shift perturbation calculations were performed using the following equation:

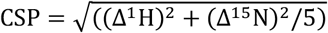

in which Δ denotes the difference in ppm of the chemical shift values of the cross-peaks between different Ub proteins.

For the chemical exchange saturation transfer (CEST) measurement,^15^N-pseudo-3D CEST experiments were collected at 318K using the established pulse sequences (36). Each experiment was conducted with a total exchange time of 400 ms, a relaxation delay of 1 s, and a *B*_1_ saturation field of 12.5 Hz or 25 Hz, covering the ^15^N range of 107 to 134 ppm. The peak intensities at the two different B_1_ fields were globally fit using ChemEx software (https://github.com/gbouvignies/ChemEx). The ^15^N CEST profiles were plotted as I/I_0_ against the B_1_ field, with the I_0_ value taken from the first transient of the acquisition.

To determine the temperature coefficients of amide protons, ^1^H, ^15^N-HSQC spectra were recorded at 4-degree intervals from 286 K to 318 K for each Ub protein. For each spectrum, ^1^H chemical shifts were referenced to the methyl proton signal of DSS. Signals were assigned by using Poky (62). Amide proton chemical shift (CS) temperature coefficients were identified as the slope obtained with a linear fitting of the CS over temperature. For the cross peaks that disappeared at one or more temperatures, the corresponding data point was omitted.

Phosphorylation of Ub proteins was conducted in the NMR buffer supplemented with 10 mM MgCl_2_, 10 mM ATP and 5 mM DTT at 298 K. The concentrations of Ub and *Ph*PINK1 were 108 μM and 0.4 μM, respectively. BEST-HSQC experiments (64, 65) were carried out to monitor the phosphorylation rates in real time with eight scans, 128 increments, and a relaxation delay of 400 ms (∼9 min for a single measurement) continuously.

The affinity of UBA binding to Ub samples was assessed by fitting the observed CSPs in the individual amino acid residues measured in the course of titration, assuming a 1:1 stoichiometry. For titration of ^15^N-labeled UBA with Ub wildtype or Ub L73A, the initial NMR sampled was prepared as 100 *μ*M in 20 mM HEPES buffer containing 150 mM NaCl at pH 7.4. A series of ^1^H,^15^N-HSQC spectra were recorded for the ^15^N-labeled UBA sampled with Ub wildtype at 26.30 *μ*M, 52.31 *μ*M, 78.03 *μ*M, 103.48 *μ*M, 128.65 *μ*M, 153.55 *μ*M, 178.18 *μ*M, and 202.55 *μ*M as the finial concentration, or Ub L73A at 24.25 *μ*M, 48.24 *μ*M, 71.97 *μ*M, 95.43 *μ*M, 118.65 *μ*M, 141.61 *μ*M, 164.33 *μ*M, and 186.81 *μ*M as the final concentration. The fitting of CSPs observed from UBA to binding isotherms by BindFit (NMR 1:1 model, http://app.supramolecular.org/bindfit/) affords the respective *K*_D_ value.

### Hydrogen Deuterium Exchange Mass Spectrometry (HDX-MS)

Peptides of Ub (WT, L67A, L69A) were identified via LC-MS/MS (ThermoFisher Orbitrap Fusion). The raw data were imported into Proteome Discover 2.4 software to identify high-score peptides. For further HDX-MS analysis, 4 μL purified Ub (WT, L67A, L69A, 8 μM) was added into 16μL D_2_O buffer (50 mM HEPES and 50 mM NaCl, pH 7.5) after incubating for various HDX time points (10s, 20s, 30s, 40s, 45s, 60s, 180s and 300s) at specific temperatures (323K, 333K, 343K) in a PCR instrument. Subsequently, the deuterium exchange was quenched by mixing with 20 μL ice-cold buffer containing 3 M guanidine hydrochloride, 1% tri?uoroacetic acid. Each quenched sample was immediately injected into the LEAP Pal 3.0 HDX platform. Upon injection, samples were passed through an immobilized pepsin column. The digested peptides were captured on a C18 PepMap300 trap column (Thermo Fisher Scientific) and separated across a C18 separating column (Thermo Fisher Scientific). The detailed experiment procedure can be referred to in our previous study (66). The percentage of deuterium content and statistical significance (unpaired t-test) were analyzed and rendered by HDX Workbench software (67).

### MD simulations and estimation of the free energy perturbation upon point mutation

Alanine mutants of Ub were introduced by removing atoms from the PDB structure 1D3Z (68). The retracted state of Ub wildtype was obtained by mutating the pSer residue back to a regular serine with the phosphoryl group deleted from PDB structure 5XK4 (35). MD simulations were performed to optimize the structure using AMBER22 package (69) with ff19SB forcefield (70). The structure was solvated in a cube containing TIP3P water molecules, with at least 10 Å padding in all directions, and the system was equilibrated and energy minimized. Unrestrained MD simulations were run for at least 20 ns for each structure at a given temperature. The distance fluctuations were evaluated using the cpptraj module in AMBER. For the steered MD simulations, the backbone heavy atoms of residues 1 to 56 were constrained with a force constant of 100 kJ/mol, while a 5 kJ/mol constant force was applied to the Cβ atom of L69 (L71 for the retracted state), at a speed of 1 Å/ns, increasing in the distance to the Cβ atom of I30 from ∼7 Å to 27 Å. The steered MD simulations were repeated 20 times with different random seeds. The destabilization energy upon introducing alanine mutations was calculated using the FoldX plugin (71) in the YASARA software at pH 7.0, 298K with 200 mM salt. For each mutant, 10 different conformers from the unrestrained MD simulations were used, and the averaged value was reported.

## Acknowledgments

This work has been supported by funds from the National Key R&D Program of China (2018YFA0507700) and the National Natural Science Foundation of China (22161132013). All NMR experiments were carried out at the Beijing NMR Center and the NMR facility of the National Center for Protein Sciences at Peking University, and the BioNMR facility, Tsinghua University Branch of China National Center for Protein Sciences (Beijing). We thank Dr. Ning Xu for his assistance in NMR data collection.

