## Supplemental Figures and Tables for "Enhancement of Protein Stability by Quenching Millisecond Conformational Dynamics"

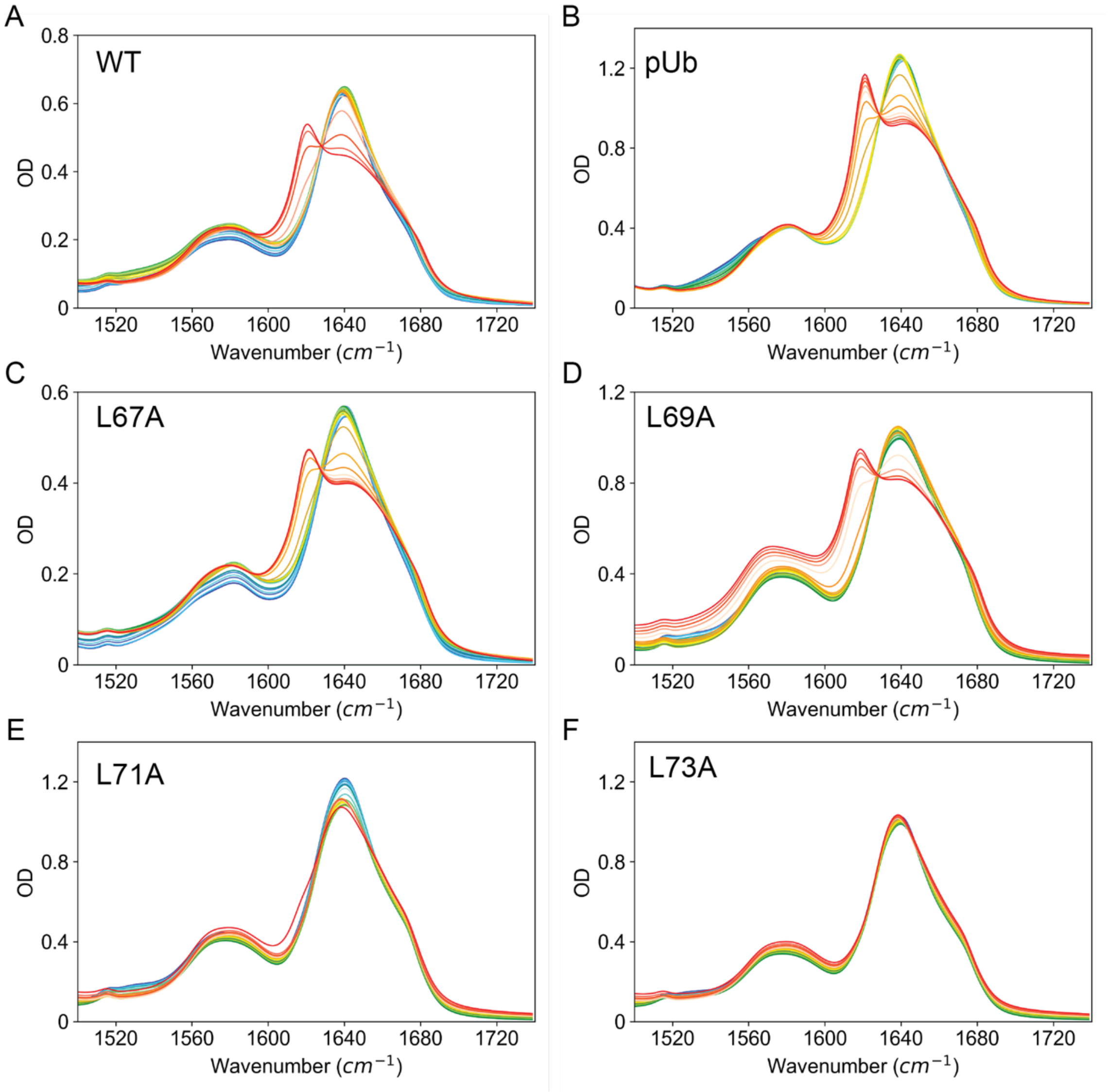


**Fig. S1.** FTIR absorption spectra of Ub proteins at increasing temperatures. Experiments were carried out at various temperatures from 30 ℃ (blue) to 99 ℃ (red).

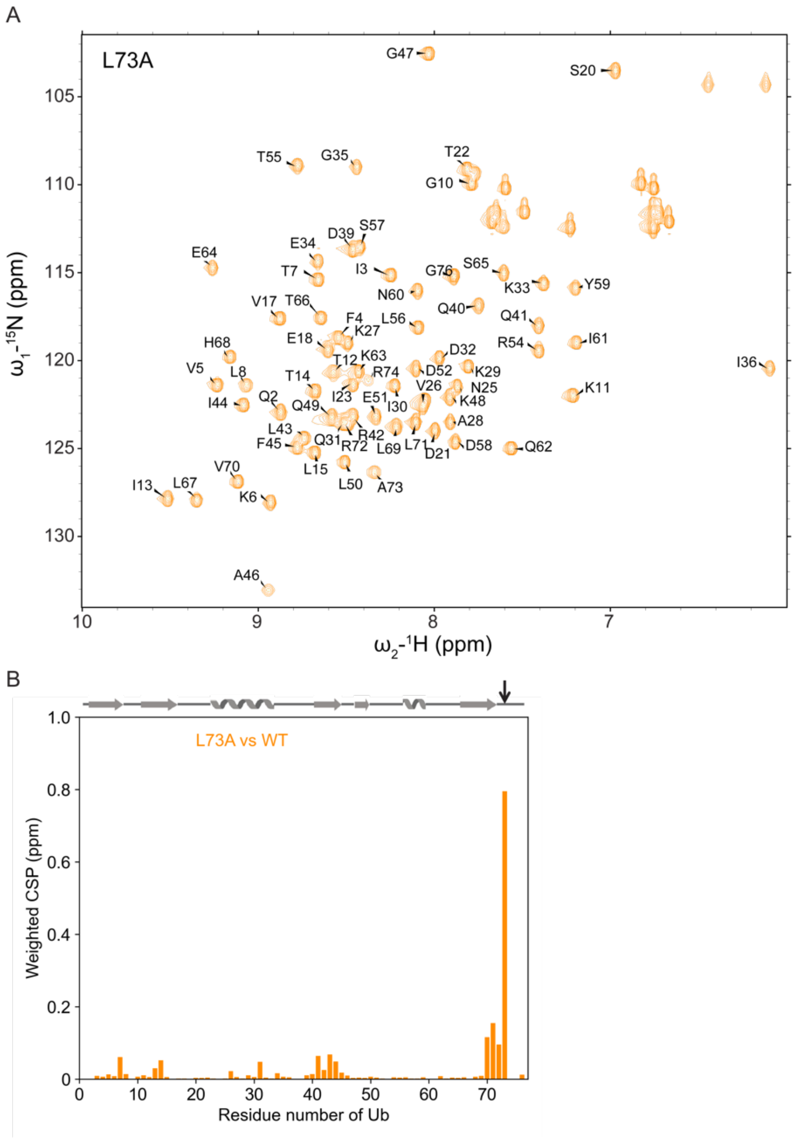


**Fig. S2.** NMR analysis of Ub L73A mutant. (A) $^{1}$H,$^{15}$N-HSQC spectrum of Ub L73A mutant. The cross-peak assignments are indicated with a one-letter amino acid code and residue number. (B) CSPs between UbL73A and Ub wildtype. Ub secondary structure and mutation site are indicated at the top.

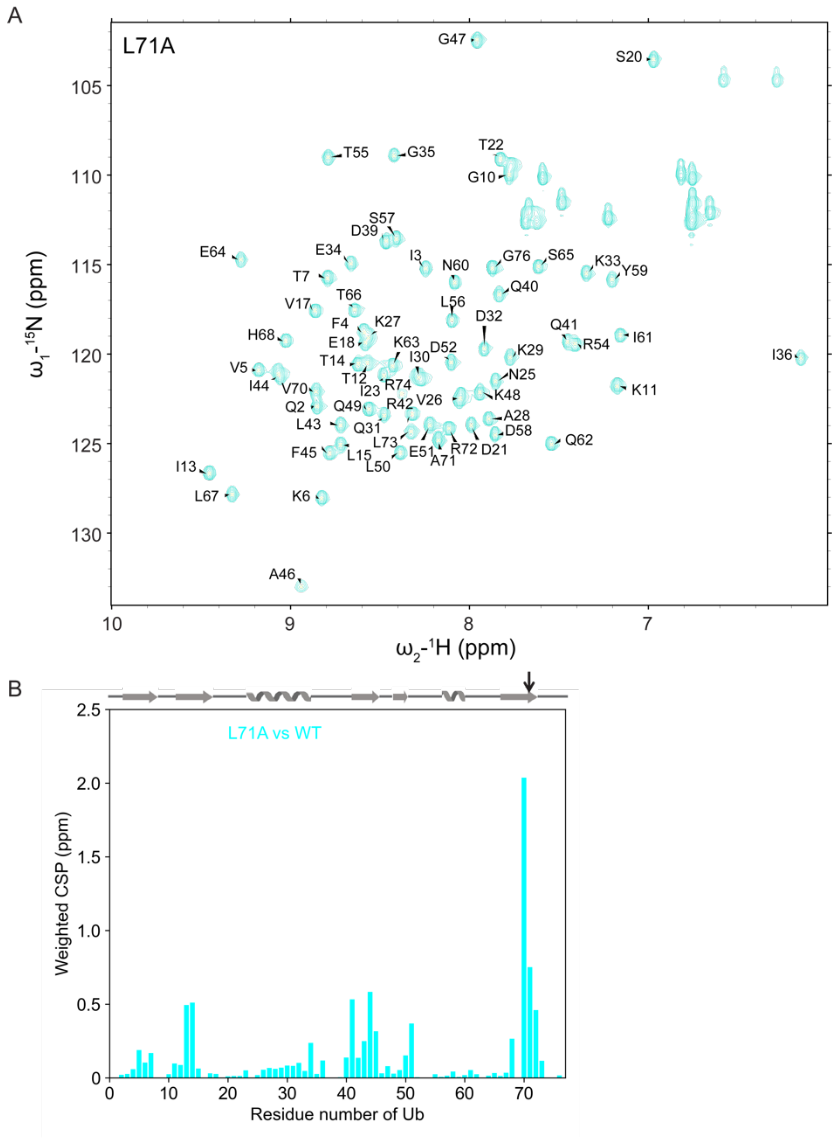


**Fig. S3.** NMR analysis of Ub L71A mutant. (A) $^{1}H,^{15}N$-HSQC spectrum of Ub L71A mutant. The cross-peak assignments are indicated with a one-letter amino acid code and residue number. (B) CSPs between UbL71A and Ub wildtype. Ub secondary structure and mutation site are indicated at the top.

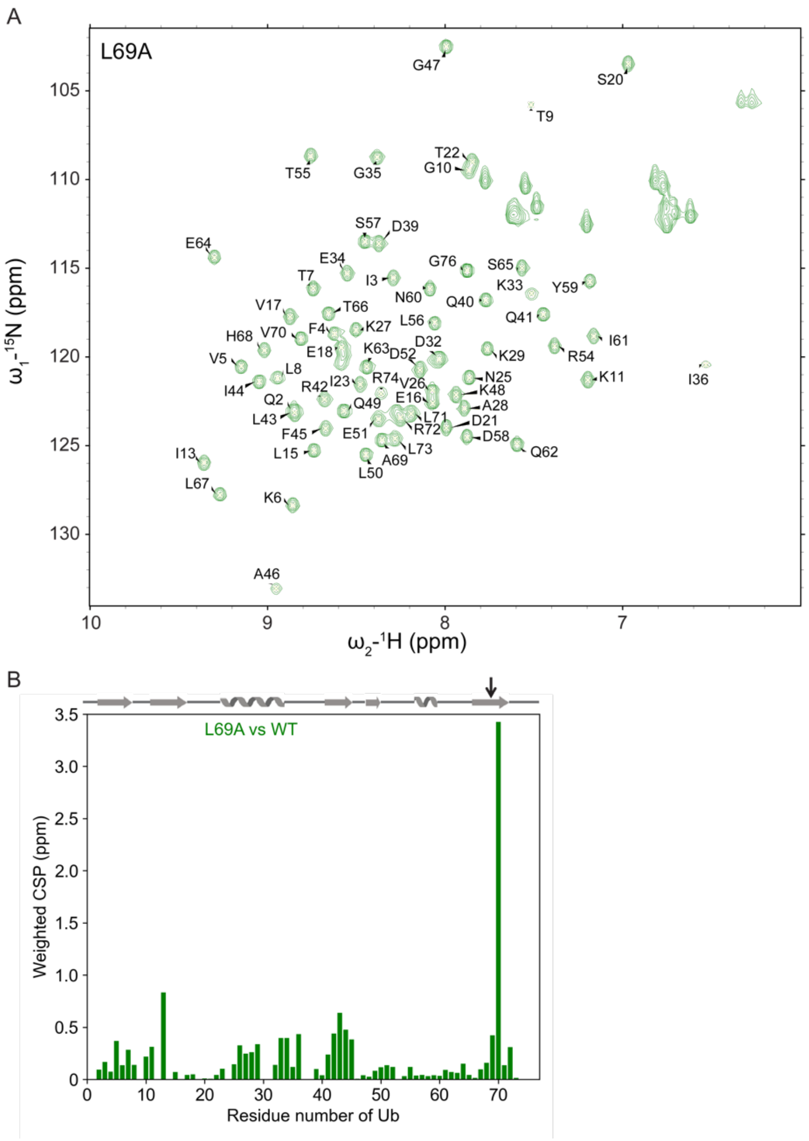


**Fig. S4.** NMR analysis of Ub L69A mutant. (A) $^{1}H,^{15}N$-HSQC spectrum of Ub L69A mutant. The cross-peak assignments are indicated with a one-letter amino acid code and residue number. (B) CSPs between UbL69A and Ub wildtype. Ub secondary structure and mutation site are indicated at the top.

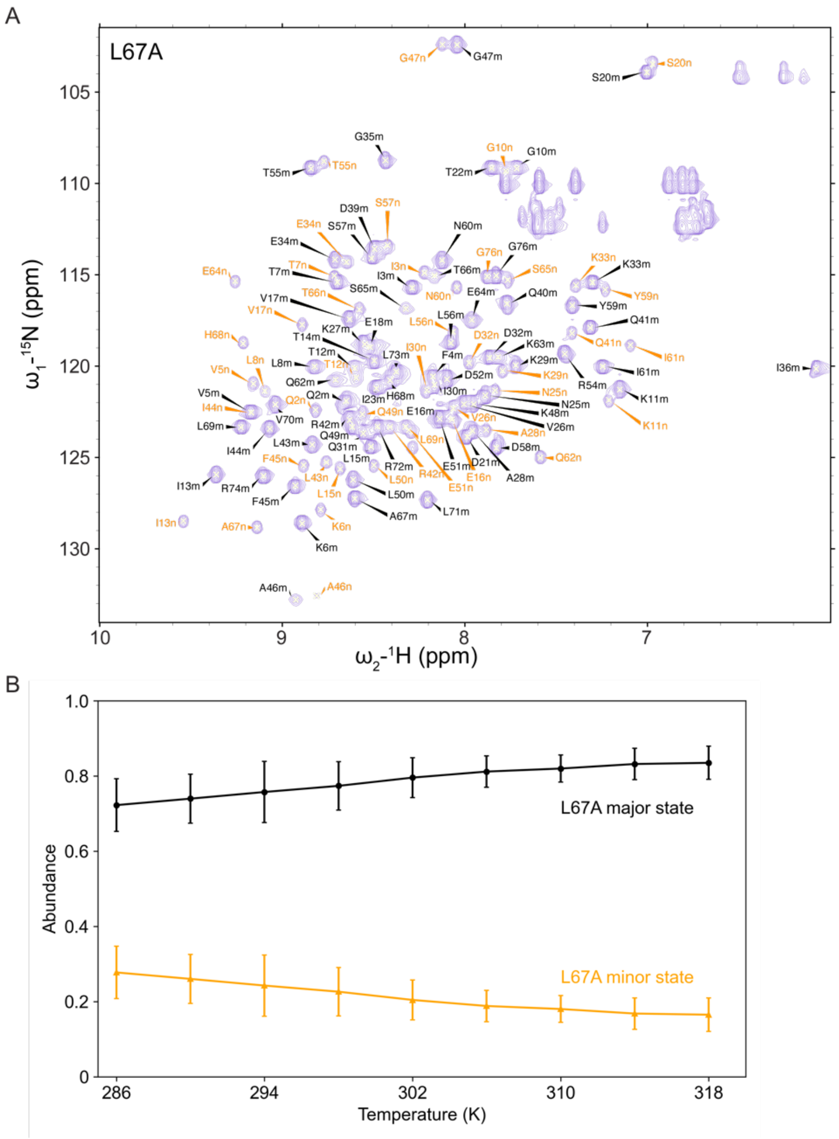


**Fig. S5.** NMR analysis of Ub L67A mutant. (A) $^{1}H,^{15}N$-HSQC spectrum of Ub L67A mutant. Chemical shift positions of the major (m, black labels) and the minor (n, orange labels) states of Ub L67A are labeled. (B) The relative abundance of the major (black) or minor (orange) state, which are approximated using peak intensities, was plotted as a function of the temperature. Data from 28 individual resonances were averaged with error bars indicating standard deviation from the average.

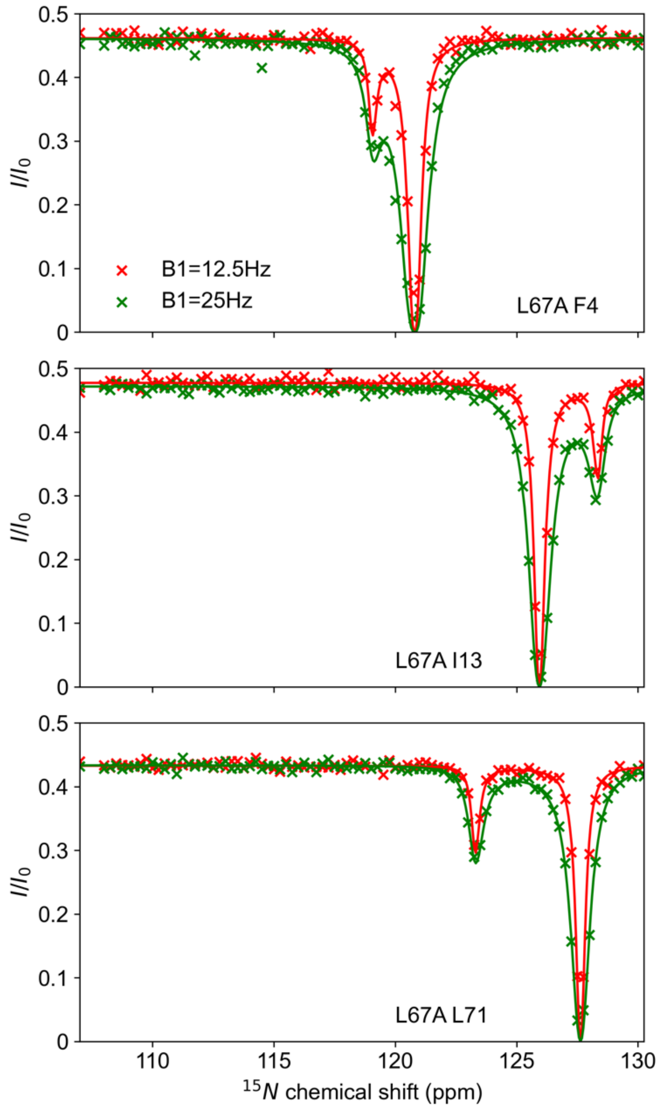


**Fig. S6.** Representative peak fits for CEST NMR experiments for Ub L67A mutant at two different $B_1$ fields collected at 318 K on an 800 MHz NMR instrument. The exchange between major and minor species is determined at 29.6±7.0 $s^{-1}$, and the minor species (relaxed state) is determined at a population of 3.8 %.

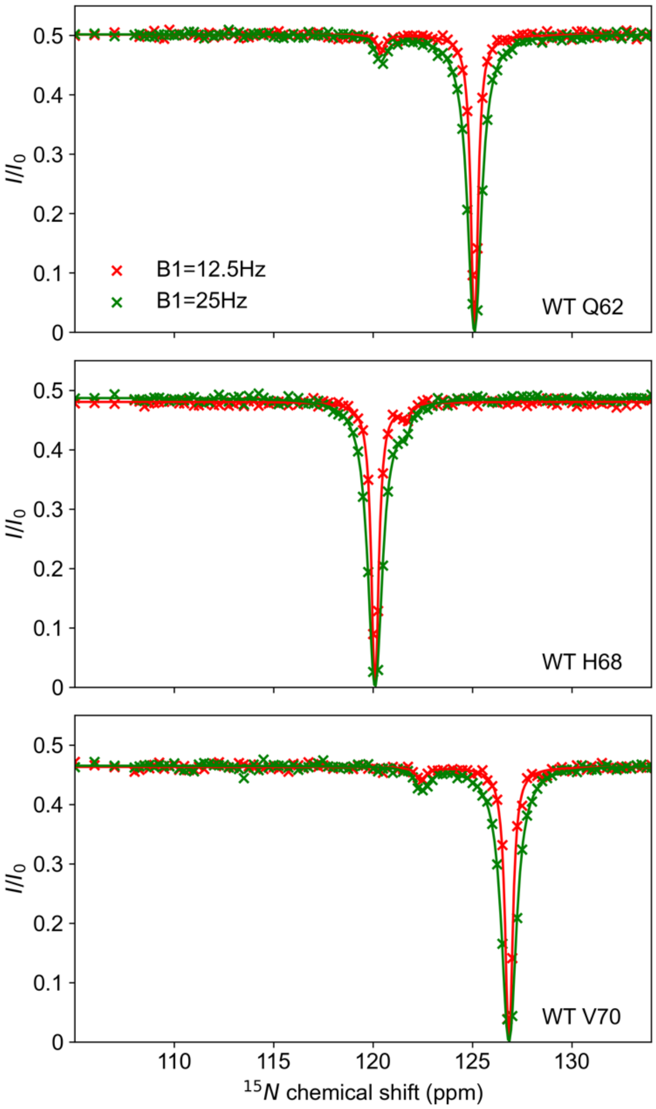


**Fig. S7.** Representative peak fits for CEST NMR experiments for the Ub wildtype at two different $B_1$ fields collected at 318 K on an 800 MHz NMR instrument. The exchange between major and minor species is determined at 55.1±14.1 $s^{-1}$, and the minor species (retracted state) is determined at a population of 0.46 %.

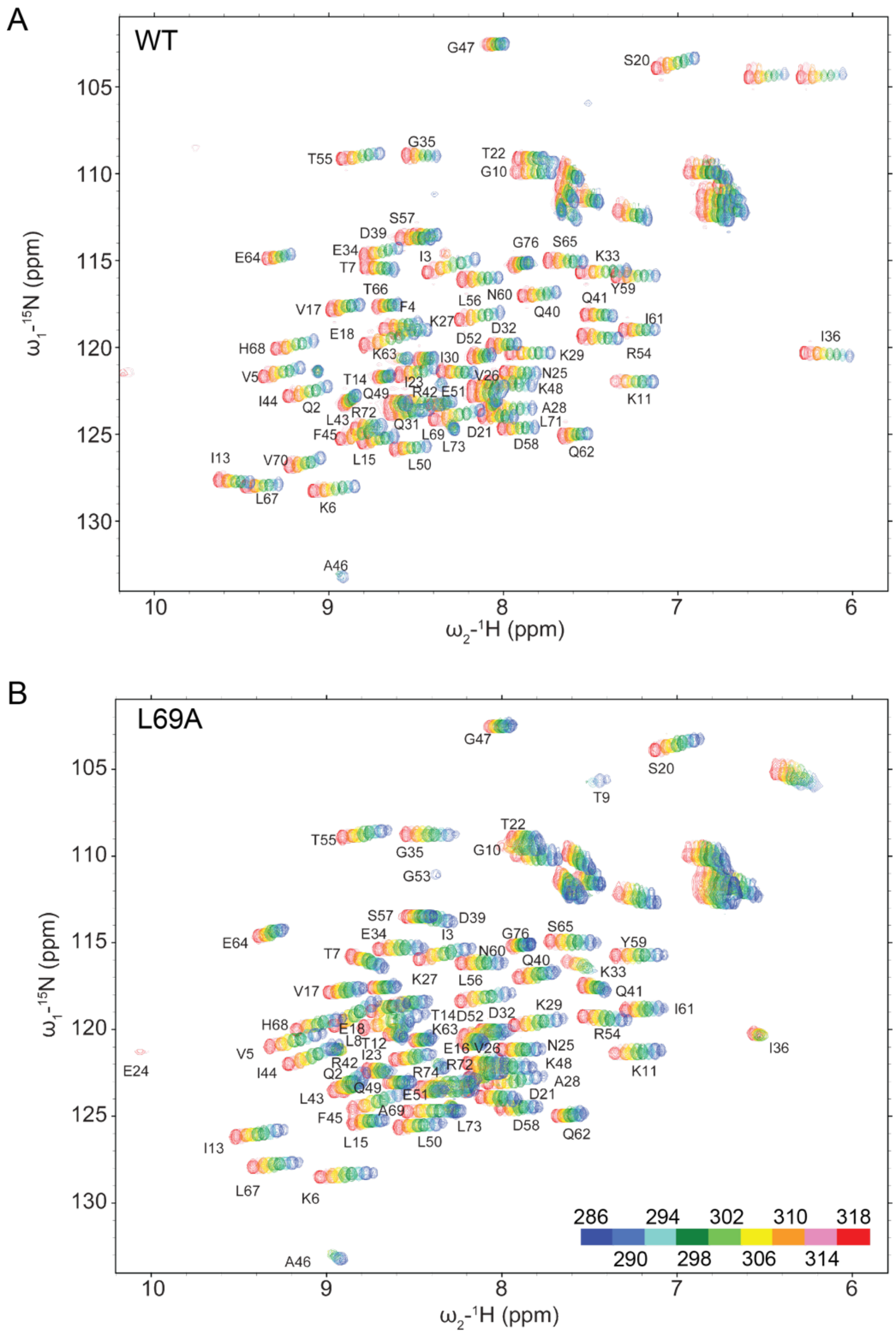


**Fig. S8.** $^{1}$H, $^{15}$N-HSQC spectra of Ub wildtype (A) and Ub L69A (B) measured at temperatures ranging from 286 K to 318 K. The temperatures are color-coded, as shown in panel B.

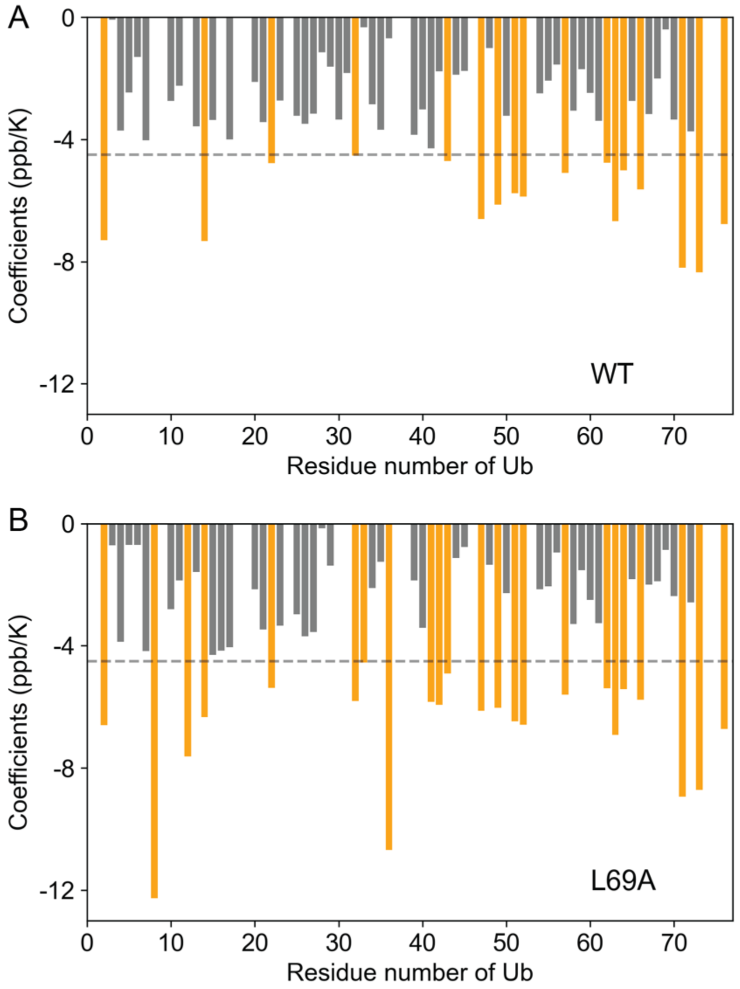


**Fig. S9.** Temperature coefficients of Ub wildtype (A) and Ub L69A (B). Residues with the temperature coefficient of more negative than -4.5 ppb/K (indicated with a dashed line) are colored orange.

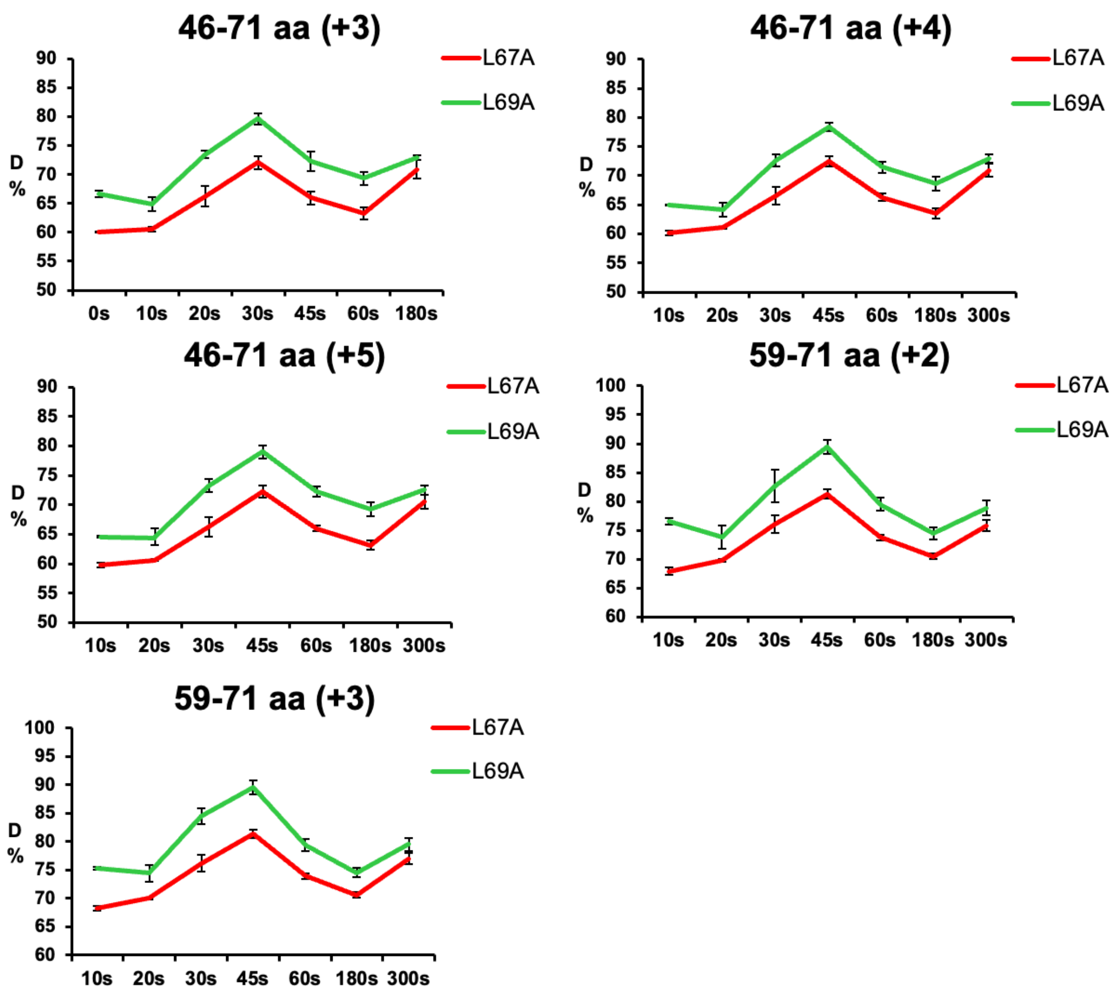


**Fig. S10.** The deuterium uptake graphs of peptides originating from Ub C-terminal peptides were shown for the L67A (red) and L69A (green) mutants.

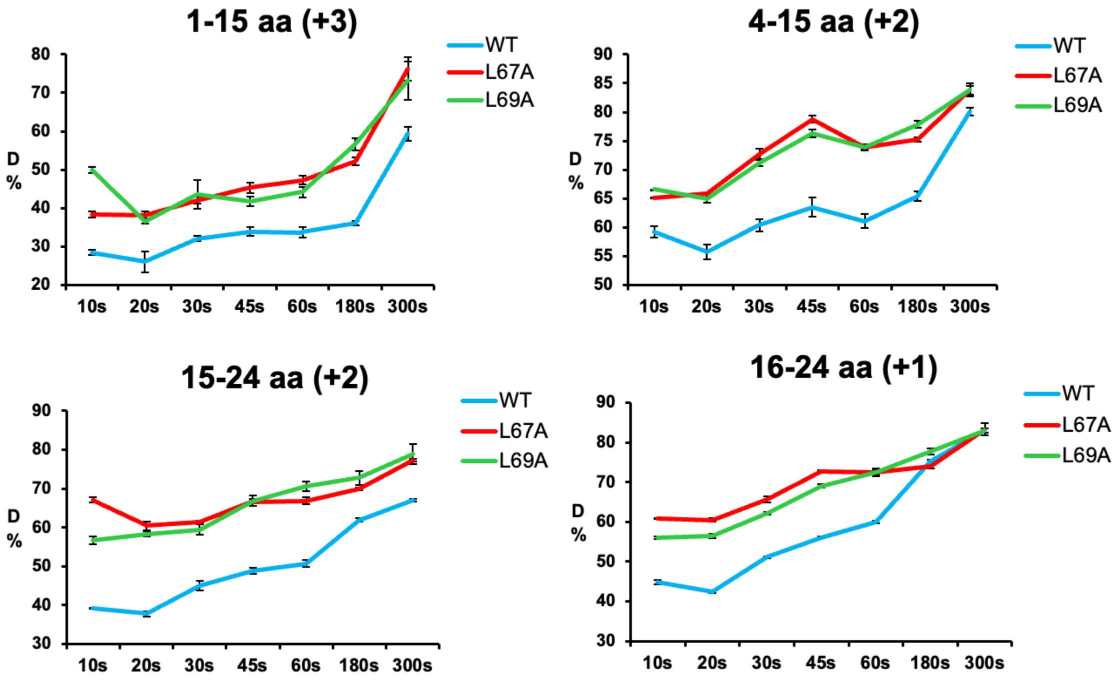


**Fig. S11.** The deuterium uptake graphs of peptides originating from N-terminal peptides were shown for wildtype (blue), L67A (red) and L69A (green) Ub proteins.

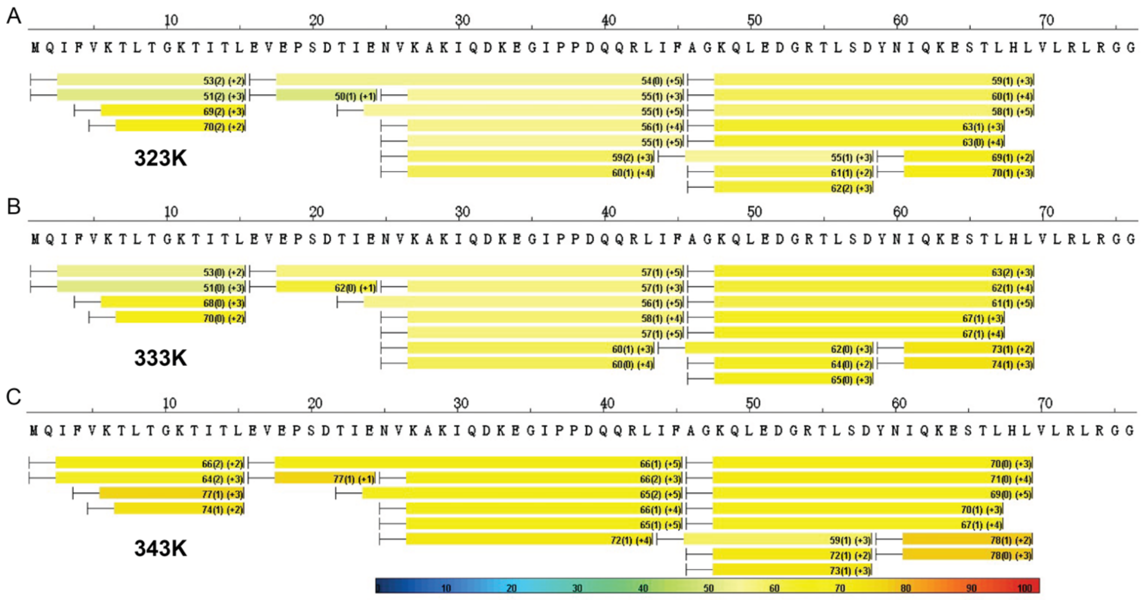


**Fig. S12.** Increase in temperature promotes structural opening and solvent exchange of Ub. Heat maps of HDX-MS analyses for Ub wildtype at 323K (A), 333 K (B) and 343K (C) Ub upon $D_2O$ exchange for 30 s. The level of deuterium substitution ratio of the labile protons is color-coded, as indicated by the legend at the bottom.

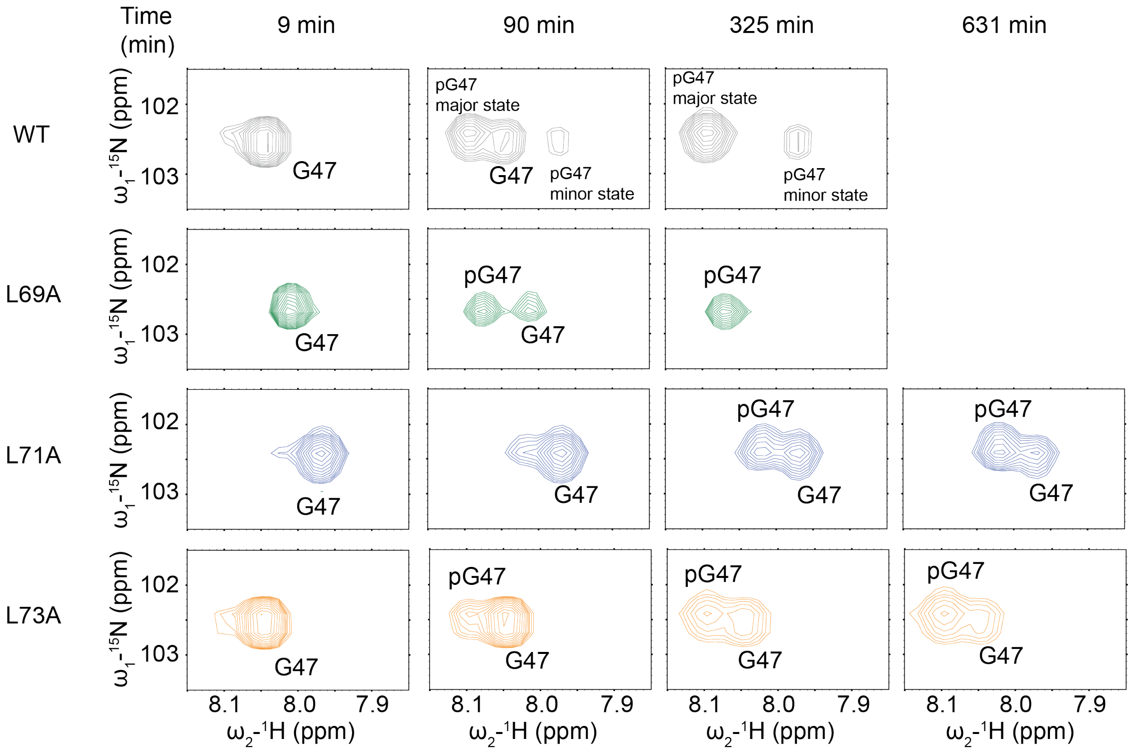


**Fig. S13.** Phosphorylation of Ub proteins by PINK1 kinase as monitored by real-time NMR. The concentrations of Ub and PINK1 kinase were 108 μM and 0.4 μM, respectively. The phosphorylation reaction was performed in an NMR tube and was monitored by a series of 2D HSQC spectra. Upon phosphorylation, Ub residues show up at different chemical shifts, which peak intensities report the progress of the reaction. The L71A and L73A mutants are much less susceptible to PINK1 phosphorylation in comparison to wildtype Ub and L69A mutant.

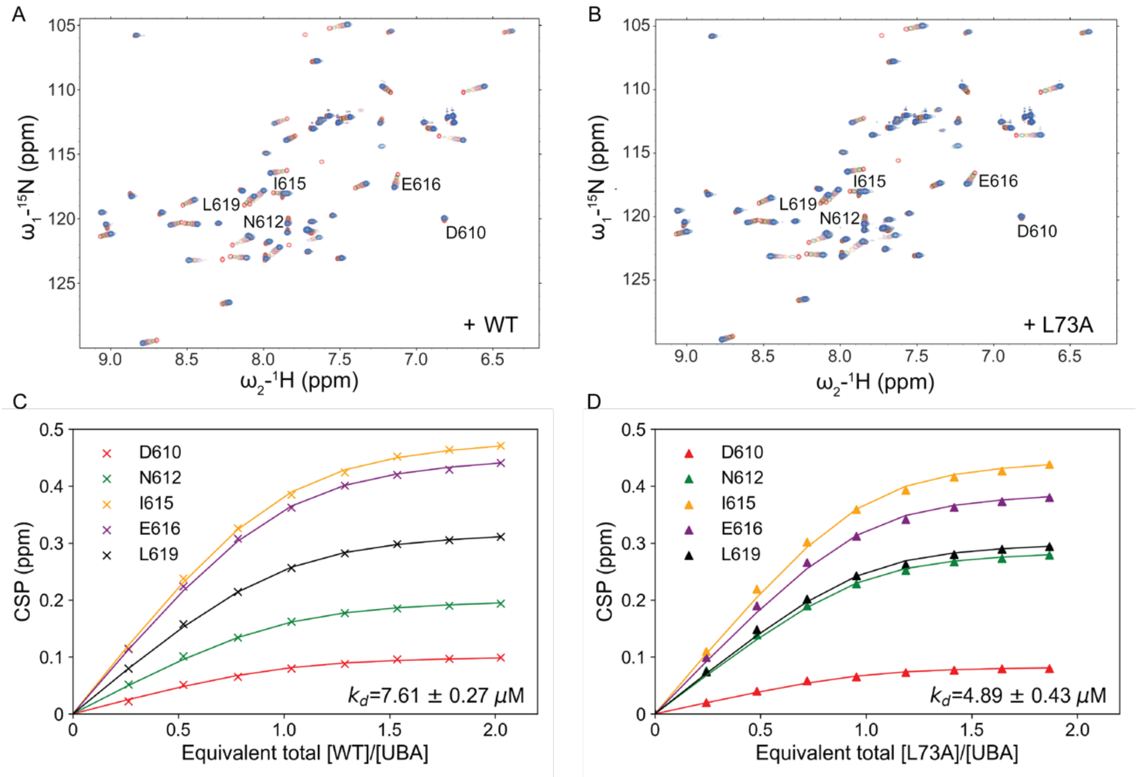


**Fig. S14.** Titration of Ub proteins with UBA domain derived from ubiquilin-2 (residues 578-621). Fitting of CSPs observed from ubiquilin-2 to binding isotherms by BindFit (NMR 1:1 model, http://app.supramolecular.org/bindfit/) affords the respective $K_d$ value.

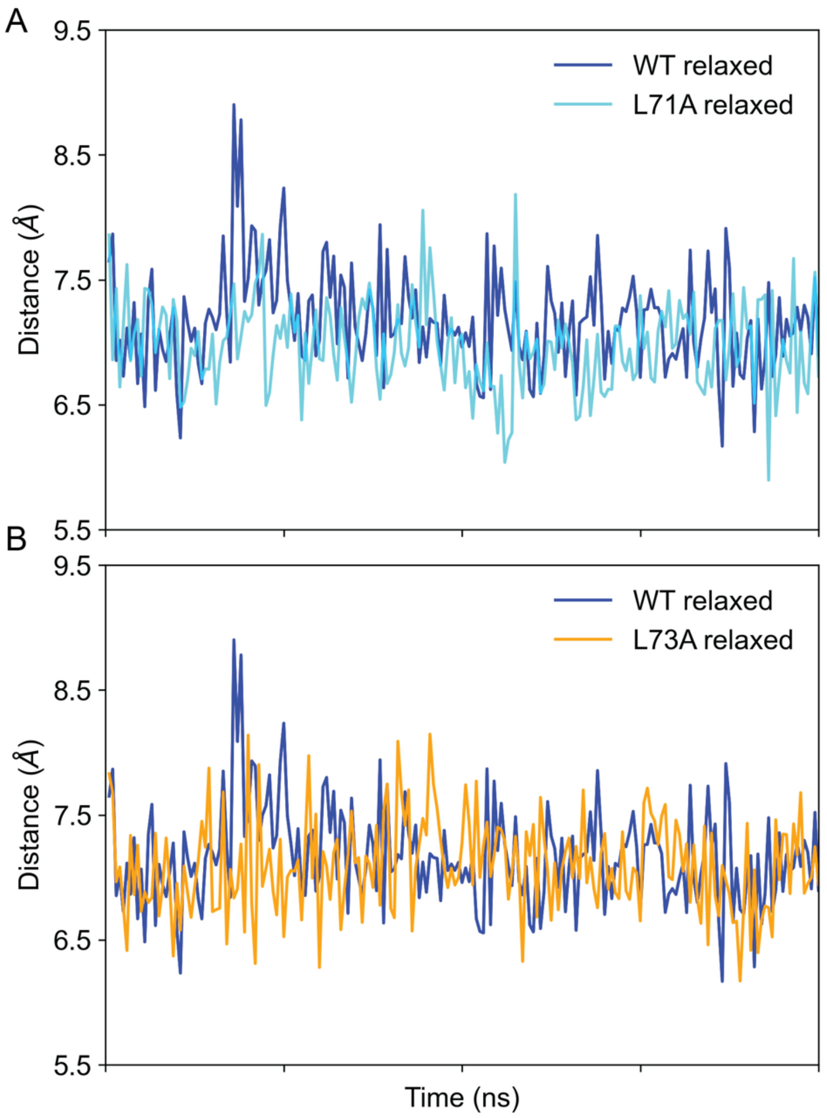


**Figure S15.** Unrestrained MD simulations of Ub wildtype, L71A mutant (A), and L73A mutant (B) in the relaxed state at 318K. A representative 20-ns trajectory is shown, and the Cβ distance between residues I30 and L69 is evaluated. The distances are 7.15±0.37 Å, 6.97±0.35 Å, and 7.09±0.38 Å, for wildtype, L71A and L73A, respectively.

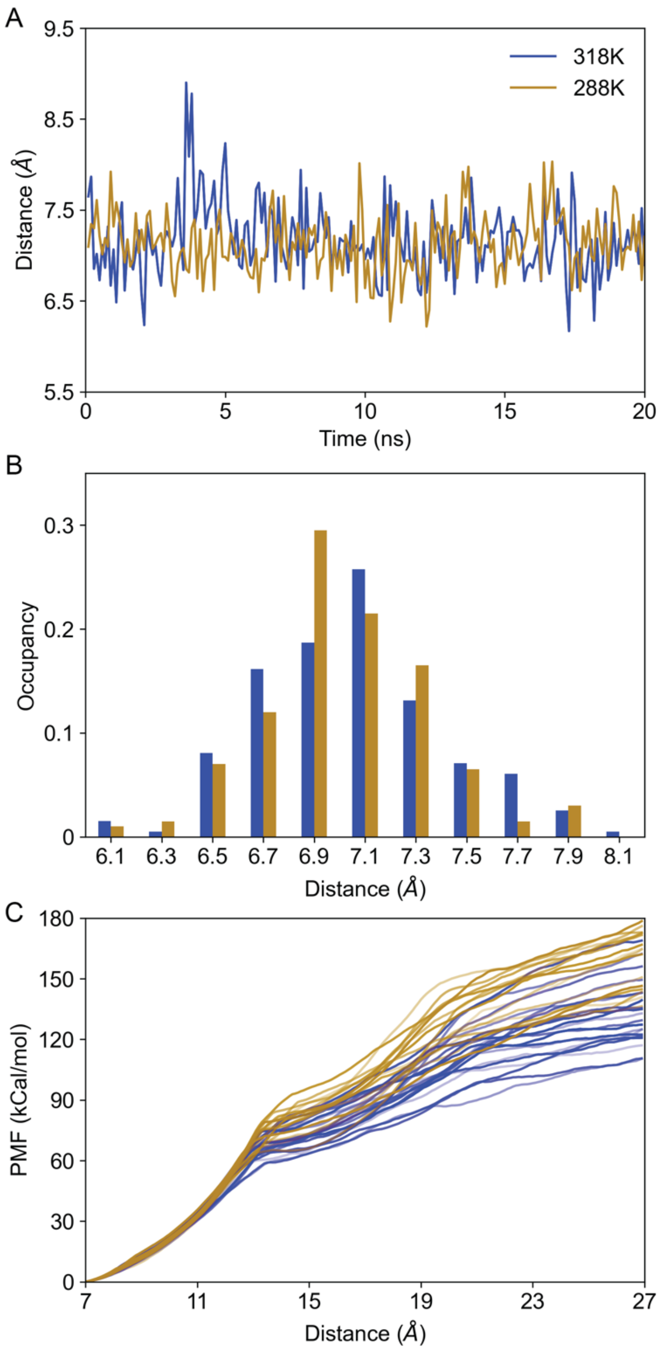


**Figure S16.** Computational analysis of Ub wildtype at 318K and 288K in the relaxed state. (A) Unrestrained MD simulations. A representative 20-ns trajectory is shown, and the Cβ distance between residues I30 and L69 is evaluated, which are 7.15±0.37 Å and 7.12 ± 0.32 Å, respectively. (B) Detailed analysis of the distance in (A), with the occurrence in each distance range at 0.2 Å interval tabulated. (C) Steered MD simulations. A constant force is applied at the L69 Cβ atom of the relaxed state. 20 independent trajectories are plotted.

**Table S1.** Cα chemical shifts for Ub wildtype and Ub L67A mutant major species.

| Residue | WT | L67A | Δ*δ* | Residue | WT | L67A | Δ*δ* |
|---|---|---|---|---|---|---|---|
| Q2 | 52.4 | 52.2 | -0.2 | Q40 | 53.04 | 52.8 | -0.24 |
| I3 | 56.93 | 56.53 | -0.4 | Q41 | 54.02 | 54.24 | 0.22 |
| F4 | 52.47 | 53.26 | 0.79 | R42 | 52.53 | 52.1 | -0.43 |
| V5 | 57.73 | 57.71 | -0.02 | L43 | 50.28 | 50.41 | 0.13 |
| K6 | 51.92 | 52.46 | 0.54 | I44 | 56.29 | 56.36 | 0.07 |
| T7 | 57.86 | 58.14 | 0.28 | F45 | 53.74 | 53.29 | -0.45 |
| L8 | 54.83 | 54.76 | -0.07 | A46 | 49.87 | 49.85 | -0.02 |
| G10 | 42.64 | 42.57 | -0.07 | G47 | 42.66 | 42.58 | -0.08 |
| K11 | 53.64 | 53.28 | -0.36 | K48 | 51.86 | 51.84 | -0.02 |
| T12 | 59.64 | 59.84 | 0.2 | Q49 | 53.24 | 53.19 | -0.05 |
| I13 | 57.43 | 57.14 | -0.29 | L50 | 51.53 | 51.79 | 0.26 |
| T14 | 59.38 | 58.66 | -0.72 | E51 | 53.31 | 52.51 | -0.8 |
| L15 | 50.03 | 50.01 | -0.02 | D52 | 53.93 | 53.41 | -0.52 |
| E16 | 52.28 | 52.3 | 0.02 | G53 | 42.42 | 42.38 | -0.04 |
| V17 | 55.84 | 55.84 | 0 | R54 | 51.61 | 51.63 | 0.02 |
| E18 | 50.06 | 50.18 | 0.12 | T55 | 57.04 | 57.03 | -0.01 |
| P19 | 62.65 | 62.58 | -0.07 | L56 | 56.07 | 56.09 | 0.02 |
| S20 | 54.72 | 54.75 | 0.03 | S57 | 58.39 | 57.97 | -0.42 |
| D21 | 53.22 | 53.16 | -0.06 | D58 | 54.86 | 54.65 | -0.21 |
| T22 | 56.98 | 57.08 | 0.1 | Y59 | 55.62 | 55.55 | -0.07 |
| I23 | 59.67 | 59.9 | 0.23 | N60 | 51.56 | 51.77 | 0.21 |
| N25 | 53.28 | 53.33 | 0.05 | I61 | 59.75 | 60.43 | 0.68 |
| V26 | 65.01 | 64.87 | -0.14 | Q62 | 50.95 | 52.27 | 1.32 |
| K27 | 56.51 | 56.45 | -0.06 | K63 | 55.14 | 55.23 | 0.09 |
| A28 | 52.76 | 52.77 | 0.01 | E64 | 55.65 | 53.51 | -2.14 |
| K29 | 57.08 | 56.99 | -0.09 | T66 | 59.82 | 58.61 | -1.21 |
| I30 | 63.5 | 63.3 | -0.2 | L67 | 51.14 | 49.42 | -1.72 |
| Q31 | 57.4 | 57.36 | -0.04 | H68 | 53.66 | 52.74 | -0.92 |
| D32 | 54.83 | 54.72 | -0.11 | L69 | 51.11 | 51.72 | 0.61 |
| K33 | 55.57 | 55.3 | -0.27 | V70 | 57.93 | 59.33 | 1.4 |
| E34 | 52.75 | 52.77 | 0.02 | L71 | 51.38 | 50.88 | -0.5 |
| G35 | 43.35 | 43.36 | 0.01 | R72 | 53.12 | 50.93 | -2.19 |
| I36 | 55.13 | 54.89 | -0.24 | L73 | 52.12 | 50.81 | -1.31 |
| D39 | 53.11 | 53.12 | 0.01 | G76 | 43.35 | 43.29 | -0.06 |
